# ASB3 expression aggravates inflammatory bowel disease by targeting TRAF6 protein stability and affecting the intestinal microbiota

**DOI:** 10.1101/2023.07.30.551192

**Authors:** Mingyang Cheng, Bin Xu, Yu Sun, Junhong Wang, Yiyuan Lu, Chunwei Shi, Tianxu Pan, Wenhui Zhao, Xiaoxu Li, Xiaomei Song, Jianzhong Wang, Nan Wang, Wentao Yang, Yanlong Jiang, Haibin Huang, Guilian Yang, Yan Zeng, Dongqin Yang, Chunfeng Wang, Xin Cao

**Author notes:** These authors contributed equally (Mingyang Cheng, Bin Xu). Correspondence: Yan Zeng, Dongqin Yang, Chunfeng Wang, Xin Cao,.

## Abstract

E3 ubiquitin ligase (E3) plays a vital role in regulating inflammatory responses by mediating ubiquitination. Previous studies have shown that ankyrin repeat and SOCS box-containing protein 3 (ASB3) is involved in immunomodulatory functions associated with cancer. However, the impact of ASB3 on the dynamic interplay of microbiota and inflammatory responses in inflammatory bowel disease (IBD) is unclear. Here we systematically identify the E3 ligase ASB3 as a facilitative regulator in the development and progression of IBD. ASB3^-/-^ mice are resistant to DSS-induced colitis. IκBα phosphorylation levels and production of proinflammatory factors IL-1β, IL-6, and TNF-α were reduced in colonic tissues of ASB3^-/-^ mice compared to WT mice. This colitis-resistant phenotype was suppressed after coprophagic microbial transfer and reversed after combined antibiotic clearance of the microbiota. Mechanistically, ASB3 specifically catalyzes K48-linked polyubiquitination of TRAF6 in IECs. In contrast, in ASB3-deficient organoids, the integrity of the TRAF6 protein is shielded, consequently decelerating the onset of intestinal inflammation. Taken together, Our findings demonstrates that ASB3 is associated with dysregulation of the colitis microbiota and promotes proinflammatory factors production by disrupting the TRAF6 stability. Strategies to limit the protein level of ASB3 in intestinal epithelial cells may help in the treatment of colitis.

## 1. Introduction

Inflammatory bowel disease (IBD) is a heterogeneous group of chronic inflammatory diseases, including Crohn’s disease (CD) and ulcerative colitis (UC), which is usually triggered by multiple complex factors.^1^ During IBD, various immune cell populations in the intestinal mucosa promote an inflammatory response by secreting proinflammatory factors. Meanwhile, intestinal epithelial cells (IECs) play an essential role in regulating intestinal immune homeostasis by maintaining the integrity of the intestinal mucosal barrier and regulating the functions of intestinal immune cells.^2^ IBD-associated genetic defects, epithelial barrier defects, dysregulated immune response, diet, and antibiotic use, among other factors, lead to early dysbiosis of the gut lamina propria (LP), which may precede the development of clinically overt disease.^3^ Continued inflammation may lead to late dysbiosis, manifested by an overall reduction in microbial diversity and loss of beneficial symbionts, leading to further exacerbation of the inflammatory response.^4^ Currently, the treatment of IBD is limited to immunotherapy. Therefore, understanding the host gene-microbiota causality and interactions in colitis provides new insights for developing therapeutic regimens with synergistic microbiota.^5^

Previous studies have shown that the E3 ubiquitin ligase, one of the critical regulators of autophagy and apoptosis, plays an essential role in normal development, tumor progression, and tissue homeostasis.^6^ E3 ubiquitin ligases promote protein degradation through the ubiquitin-proteasome system (UPS). One class of E3 ubiquitin ligases is called SOCS family proteins because of the SOCS box domain.^7^ Ankyrin (ANK) repeat and SOCS box containing protein 3 (ASB3) is a member of the ASB family. This family has 18 members, and the protein structure includes the N-terminal ankyrin repeat and the C-terminal SOCS cassette.^8^ The N-terminal domain of the ASB3 protein is responsible for substrate recognition, and the C-terminal SOCS box performs E3 enzyme function.

Ubiquitination modification is a crucial posttranslational modification (PTM) in the inflammatory response, affecting the function of essential proteins of the signaling pathway.^9^ Dysregulation of E3 ubiquitin ligase-related response processes has been reported in IBD and numerous immune-related diseases.^10^ For instance, ASB1 promotes TAB2 stability by inhibiting K48-linked polyubiquitination, which leads to enhanced NF-κB and MAPK activation.^11^ MARCH3 negatively regulates IL-6/STAT3 signaling to limit colitis by targeting IL-6Rα ubiquitination of IL-6Rα at K401 and subsequent degradation.^12^ In addition, TRIM34 regulates intestinal inflammation by controlling Muc2 cytokinesis in colonic goblet cells (GCs).^13^ Thus, more elucidation of the functions and targets of E3 ubiquitin ligases will provide additional avenues for the treatment of infectious diseases. To date, despite new evidence of the importance of ASB3 regulation in cancer development, its role in colonic inflammation has not been evaluated.

Here, we investigated the physiological role of ASB3 in the intestine and found that ASB3-deficient mice exhibited reduced susceptibility to dextran sodium sulfate (DSS)-induced colitis. Further experiments showed that ASB3 deficiency directly protects against Bacteroidetes and inhibits *Lactobacillus* overgrowth, thus favoring the growth of *Akkermansia*. Moreover, we demonstrate that IECs ASB3 modulates proinflammatory factors by specifically enhancing K48-linked polyubiquitination of TRAF6. This novel mechanism precedes the dysregulation of gut ecology and reinforces the role of ASB3 in maintaining inflammatory signaling and a healthy gut microbial ecology.

## 2. Results

### 2.1. ASB3 expression is upregulated in IBD patients and in mice with DSS-induced colitis

To explore the potential role of ASB3 in IBD, we examined the expression levels of ASB3 in the colonic tissues of UC and CD patients. The results showed that the mRNA expression of ASB3 was upregulated in both UC (n=5) and CD (n=5) compared to controls (Figure 1a, c). Consistent with the transcriptional level, the expression of ASB3 protein was enhanced at the site of IBD inflammation and was accompanied by damage to the intestinal barrier (Figure 1b, d). We then verified the level of ASB3 in the colons of mice. There was an increase in IκBα phosphorylation and in ASB3 expression in the colonic tissue from DSS-treated FVB mice (Figure 1e, f). Furthermore, similar results were observed in colon organoids and HT-29 cells stimulated with TNF-α (Figure 1g-i), suggesting that intestinal inflammation is accompanied by abnormally high ASB3 expression. We also observed that ASB3 expression levels were higher in the colonic mucosa of IBD patients than in the colonic mucosa of uninflamed donors (Figure 1j). In conclusion, abnormally high expression of ASB3 was positively correlated with inflammation in IBD tissues.

**Figure. 1.**
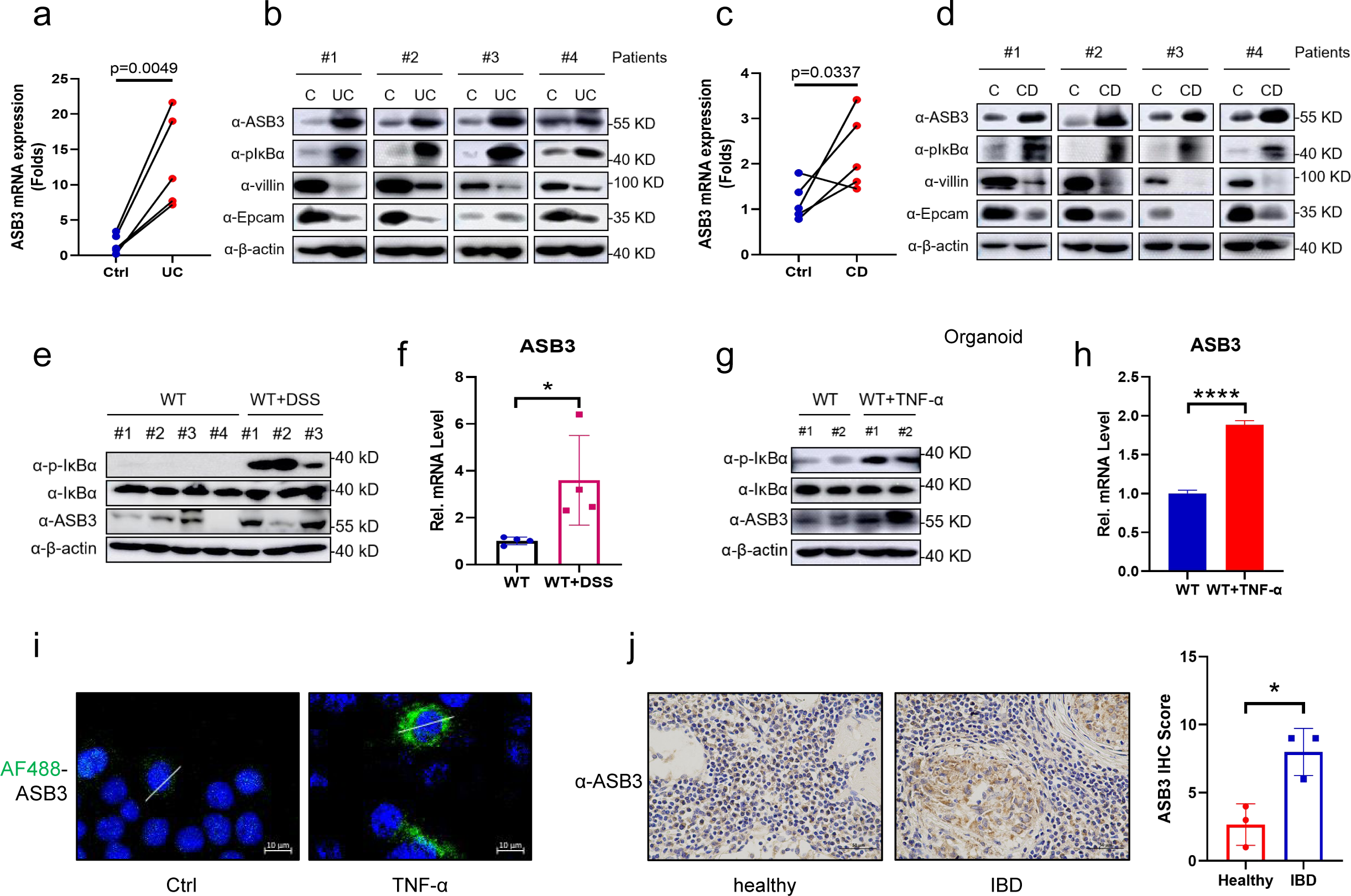
Abnormal ASB3 expression levels in IBD patients and in mice with DSS-induced colitis. **a** The mRNA expression of ASB3, IL-1β, IL-6, and TNF-α was measured in both UC groups (n=5) relative to controls. **b** Representative immunoblotting of p-IκBα, IκBα, ASB3, villin, EpCAM and β-actin protein expression in UC. **c** The mRNA expression of ASB3, IL-1β, IL-6, and TNF-α was measured in both CD groups (n=5) relative to controls. **d** Representative immunoblotting of p-IκBα, IκBα, ASB3, villin, EpCAM and β-actin protein expression in CD. **e** The expression of p-IκBα, IκBα, ASB3, and β-actin proteins in the colons of DSS-treated or untreated mice was detected by Western blotting. **f** The mRNA expression levels of ASB3 in the colons of DSS-treated or untreated mice were determined by qPCR assay. **g** The expression of p-IκBα, IκBα, ASB3, and β-actin proteins in TNF-α-treated or untreated organoids was detected by Western blotting. **h** The mRNA expression levels of ASB3 in the organoids of TNF-α-treated or untreated mice were determined by qPCR assay. **i** HT-29 cells were treated with TNF-α (150 ng/ml) for 12 h, then stained with indicated antibody and secondary antibody. The nuclei were stained by DAPI. Scale bars, 50 µm. **j** Representative IHC staining (left) and quantification (right) of ASB3 in colon tissues collected from IBD patients. Scale bars, 50 µm. P values less than 0.05 were considered statistically significant (\**P* < 0.05, \*\**P* < 0.01, \*\*\**P* < 0.001, \*\*\*\**P* < 0.0001). Differences between groups in **a**, **c**, **f**, **h** and **j** by unpaired Student’s t test.

### 2.2. ASB3^-/-^ mice are resistant to DSS-induced colitis

To further investigate the function of ASB3 in the development of colitis, ASB3^-/-^ and WT mice were stimulated with 3% DSS for 6 days and then allowed to recover with normal drinking water for 2 days. The lack of ASB3 expression in the healthy state did not lead to anatomical abnormalities or spontaneous inflammation of the intestine (Figure S1b-d, Supporting Information). Nevertheless, our results showed that ASB3^-/-^ mice had a significantly lower rate of weight loss than WT mice after DSS challenge (Figure 2a). We observed that WT mice died during the acute phase of DSS challenge, while all ASB3^-/-^ mice survived (Figure 2b). In addition, the reduced severity of colitis observed in ASB3^-/-^ mice was manifested by less diarrhea and fewer bloody stools, longer colonic length, and reduced clinical scores (Figure 2c-f). To assess this hypoinflammatory phenotype, we analyzed histopathological sections of the colon. Consistent with the clinical features described above, ASB3^-/-^ mice exhibited less severe submucosal inflammatory cell infiltration and less severe epithelial or crypt erosion than WT mice (Figure 2g). Epithelial cells include enterocytes, goblet cells, enteroendocrine cells, Paneth cells, and M cells, which are tightly connected and covered with mucus. Intestinal barrier damage and epithelial cell depletion are regarded as important characteristics associated with mucosal inflammation.^14^ We further confirmed by qPCR and Alcian blue staining that ASB3 expression facilitated a reduction in the number of cupped cells (Figure 2h, i). We also found that the absence of ASB3 expression during inflammation attenuated the damage to intestinal epithelial cells (Figure 2j). Our findings suggest that ASB3 deficiency protects against DSS-induced colonic inflammation.

**Figure. 2.**
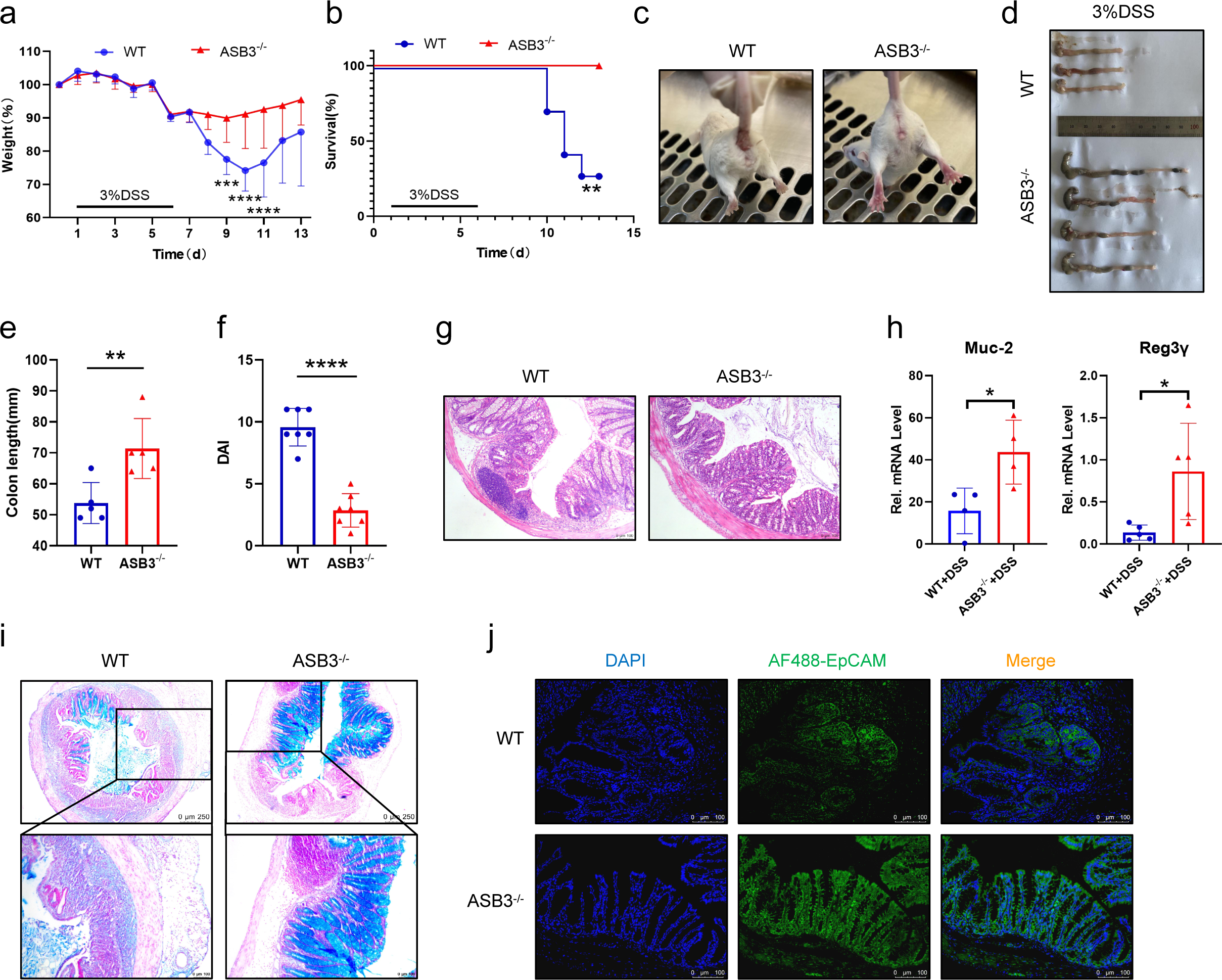
ASB3 deficiency provides protection against DSS-induced colitis. **a, b** Body weight (**a**) and survival (**b**) of conventionally raised wild-type (WT) and ASB3^-/-^ mice treated with 3% DSS (above horizontal axes) (WT, n=7; ASB3^-/-^, n=7); the results are presented relative to initial values, set as 100% (throughout). **c** Representative images of diarrhea/bloody stools observed on day 6. **d, e** Colon length on day 8. **f** DAI scores were determined on day 6. **g** Representative images of pathological H&E stained colon sections collected on day 8. Scale bar, 100 μm. **h** The mRNA expression levels of Muc-2 and Reg3γ were determined by qPCR assay. **i** Representative images of Alcian Blue staining of colon tissues of WT and ASB3^-/-^ mice. Scale bar, 250 or 100 μm. **j** IF for EpCAM in the colon; representative image of 3 mice/genotype. Scale bar, 100 μm. Differences between groups in **e**, **f**, **h** were determined by unpaired Student’s t test; in **a** by One-way ANOVA; in **b** by log-rank test.

### 2.3. ASB3 exacerbates colitis by promoting the release of proinflammatory factors

The development of IBD is closely related to intestinal cytokine storms.^15^ To determine whether ASB3 regulates the production of proinflammatory cytokines, we examined the production of IL-1β, IL-6, and TNF-α in the mouse colon.^16^ As expected, the transcript levels and secretion levels of IL-1β, IL-6, and TNF-α in the colon of ASB3^-/-^ mice after DSS challenge were much lower than those in WT mice (Figure 3a, b). Moreover, ASB3 deficiency resulted in diminished phosphorylation of NF-κB-dependent proteins, exemplified by IκBα (Figure 3c). These findings establish a link between ASB3 expression and increased IBD exacerbation and solidify that ASB3 expression may exacerbate colonic inflammation through excessive activation of immune signaling. To a great extent, mucosal immune cells directly alter the function of epithelial cells, thus adapting organ function to changing demands.^17^ The ILC population, which plays a critical regulatory role in intestinal inflammation, attracted our attention. We observed a significant decrease in the percentage of ILC3s in Lin-cells in colonic lamina propria lymphocytes (cLPLs) of WT mice but not in ASB3^-/-^ mice after administration of the drug compared to controls (Figure 3d). WT mice exhibited loss of the IL-22^+^ ILC3 subpopulation in colonic LPLs and reduced mRNA expression of IL-22 (Figure 3e-g). ILC3-derived IL-22 is essential for tissue repair.^18^ Hence, we are also more convinced by this evidence of the importance of ASB3 expression in maintaining homeostasis in the intestine.

**Figure. 3.**
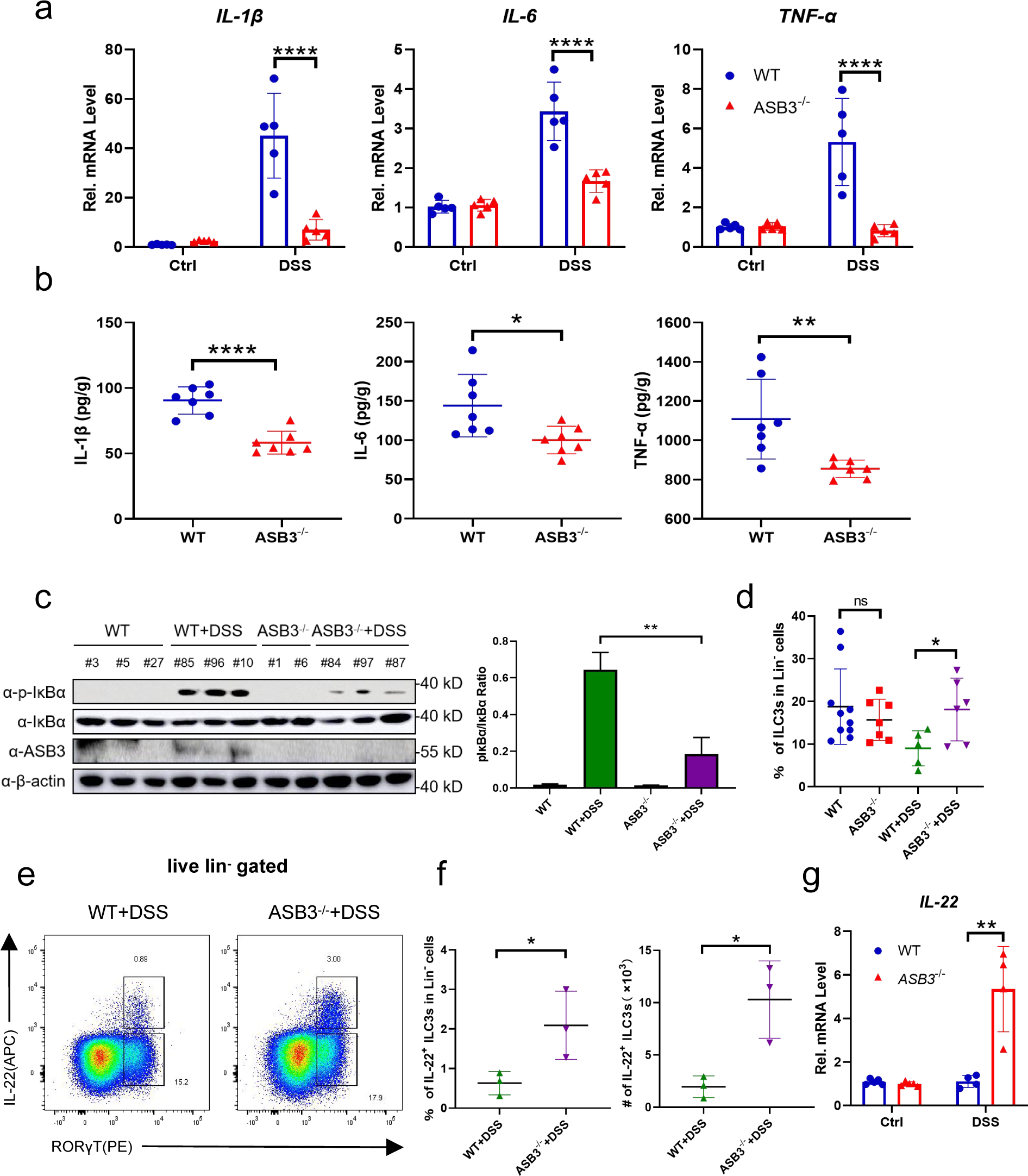
ASB3 deficiency inhibits the synthesis and release of proinflammatory cytokines **a** qPCR analysis of IL-1β, IL-6, and TNF-α mRNA expression in colon tissue sections from WT (n=5) and ASB3^-/-^ mice (n=5) treated with 3% DSS. **b** Colonic secretion of cytokines. **c** The expression of p-IκBα, IκBα, ASB3, and β-actin proteins in colon tissues from WT (n=6) and ASB3^-/-^ mice (n=5) treated with or without 3% DSS was detected by Western blotting. **d** Frequencies of ILC3s in cLPLs from WT and ASB3^-/-^ mice treated with or without 3% DSS. **e** Proportions of IL-22^+^ ILC3 cells in the colon of WT (ASB3^+/+^) and ASB3^-/-^ mice treated with 3% DSS. **f** Frequencies and absolute number of IL-22^+^ ILC3s in cLPLs from WT (n=9) and ASB3^-/-^ (n=9) mice treated with 3% DSS. **g** The mRNA expression levels of IL-22 were determined by qPCR assay. Differences were determined in **b**, **c**, **d**, **f** by unpaired Student’s t test; in **a**, **g** by One-way ANOVA.

### 2.4. Attenuated colitis in ASB3^-/-^ mice is partially dependent on altered microbiota

Currently, the etiology of IBD is thought to be caused by inappropriate interactions between host cells and the gut microbiota secondary to increased inflammation involving the overgrowth of harmful opportunistic bacteria or pathogens and the loss of beneficial bacteria.^19^ To determine whether regulation of microbiota by ASB3 resulted in changes in specific bacteria, we performed a high-throughput sequencing analysis of 16S rDNA isolated from WT and ASB3^-/-^ mice before and after DSS challenge. These bacteria were originally derived from the same ASB3^+/-^ parents in our animal facility. The petal and box plots show that the ASB3^-/-^ mice had a lower relative abundance of fecal bacteria than the WT mice (Figure S2a, b, Supporting Information). Notably, principal coordinate analysis revealed no segregation between genotypes at steady state. However, distinct colonies were formed in ASB3^-/-^ mice after DSS challenge (Figure S2c, Supporting Information). At the phylum level, the abundance of Bacteroidetes was elevated in ASB3^-/-^ mice before and after DSS treatment, and the abundance of Muribaculaceae was reduced (Figure S2d, Supporting Information). Furthermore, at the genus level, ASB3^-/-^ mice exhibited a reduced abundance of *Lactobacillus* and an increased abundance of *Akkermansia* (Figure S2e, Supporting Information). To determine whether the slowing of colitis observed in ASB3^-/-^ mice was associated with the composition of the intestinal microbiota, we performed microbiota-transfer studies by cohousing mice, which leads to exchange of the microbiota through coprophagia. Six-to eight-week-old WT and ASB3^-/-^ mice were either housed individually (SiHo mice) or cohoused (CoHo mice) for 6 weeks and then treated with 3% DSS (Figure 4a). We found that SiHo WT mice and SiHo ASB3^-/-^ mice continued to have significantly different rates of weight loss after receiving DSS (Figure 4b). However, ASB3^-/-^ mice cohoused with WT mice (CoHo ASB3^-/-^ mice) exhibited similar rates of weight loss as WT mice (CoHo WT mice) in the same littermates (Figure 4c). Notably, CoHo ASB3^-/-^ mice showed acute phase death on day 9, a trend similar to that of WT mice after either littermates or non-littermates DSS administration (Figure 4d). Furthermore, CoHo ASB3^-/-^ and their WT cage companions were similar across almost all measures, including colon length, disease activity score, and histopathological changes (Figure 4e-g). We also demonstrated that microbial exchange somewhat alleviated IκBα phosphorylation and inflammatory factor (IL-1β, IL-6, and TNF-α) expression in the colons of CoHo WT mice (Figure 4h, i). We note that while the transferred microorganisms reduced the enteritis phenotype within the same litter of WT mice, the difference was narrowed but still existed. We speculate that the role in IBD pathogenesis of inappropriate host gene-microbe interactions may be causal. To test this, we first cleared the intestinal flora using a combination of antibiotics and then induced colitis using DSS (Figure 4j). Interestingly, despite the reversal of the colitis phenotype in ASB3^-/-^ mice, the phosphorylation level of IκBα in the colon was lower than that in WT mice (Figure 4k). These data suggest that ASB3 expression may preferentially determine the substrate of intestinal inflammation, amplifying microbial signals through many inflammatory factors, thereby exacerbating intestinal barrier damage and the development of colonic inflammation.

**Figure. 4.**
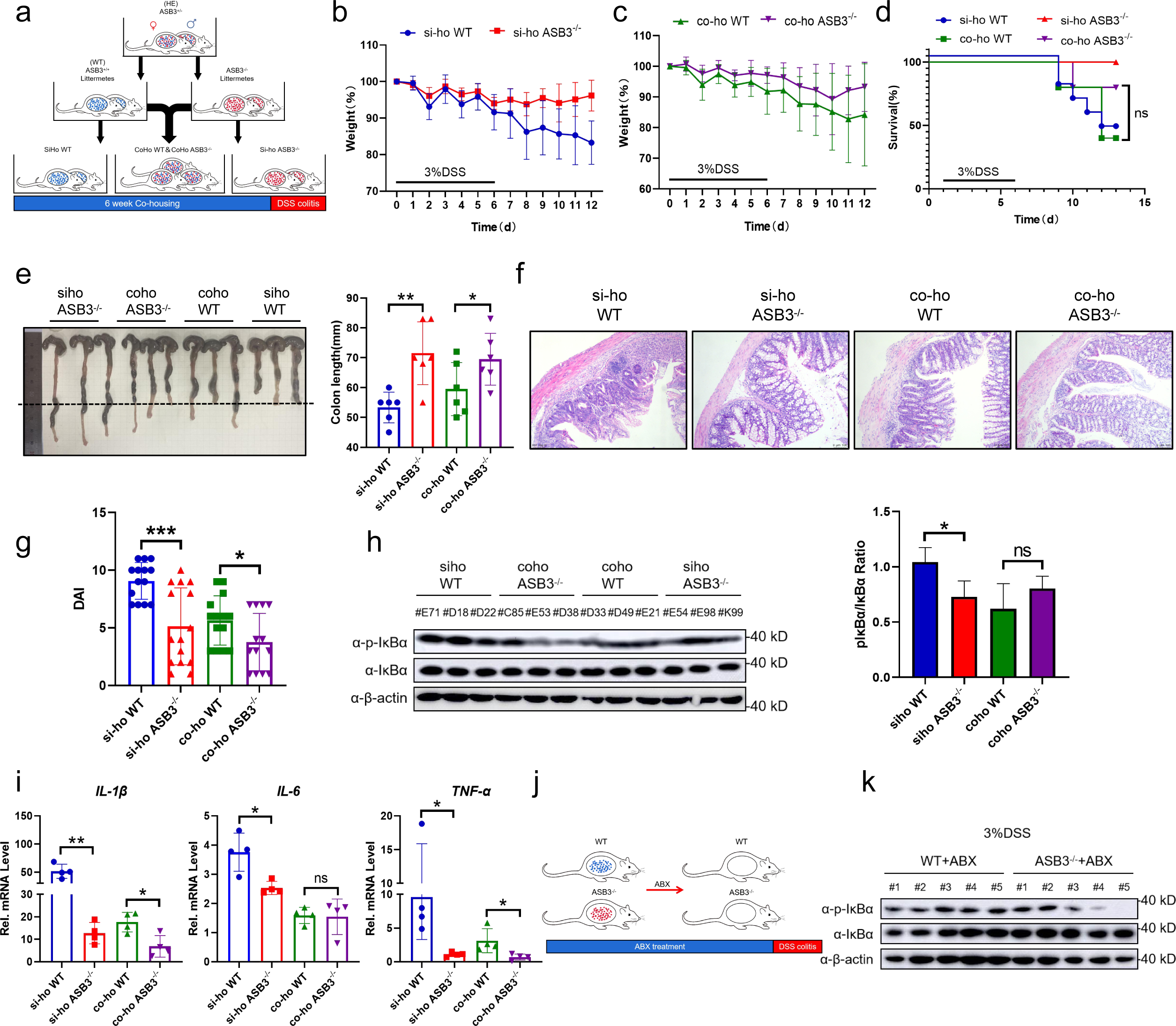
Commensal microbiota from ASB3^-/-^ mice enhances protection against DSS-induced experimental colitis in WT mice. **a** Schematic representation of the SiHo-CoHo strategy and DSS treatment. **b-d** Body weight (**b, c**) and survival (**d**) of SiHo-WT, SiHo-ASB3^-/-^, Coho-WT or Coho-ASB3^-/-^ mice during DSS-induced colitis. **e-g** Colon length (**e**) representative H&E staining (**f**) and DAI (**g**) of WT, ASB3^-/-^, Coho-WT or Coho-ASB3^-/-^ mice on day 8 after DSS induction. Scale bar, 50 µm. **h** The protein expression of p-IκBα, IκBα, and β-actin was detected by Western blotting. **i** The mRNA expression levels of IL-1β, IL-6, and TNF-α were determined by qPCR assay. **j** Schematic representation of the DSS challenge after ABX treatment. **k** The protein expression of p-IκBα, IκBα and β-actin was detected by Western blotting. Difference were determined in **e**, **g**, **h** by unpaired Student’s t test; in **d** by log-rank test.

### 2.5. ASB3 promotes NF-κB activation by interacting specifically with TRAF6

A recent study showed that overexpression of ASB17 increased the level of LPS-mediated NF-κB activation in THP-1 cells.^20^ However, the mechanism of ASB3-induced NF-κB activation is unclear. To determine the role of ASB3 in NF-κB signaling pathway activation, we stimulated WT and ASB3-overexpressing 293T cells with TNF-α and found that overexpression of ASB3 significantly upregulated the phosphorylation level of IκBα (Figure 5a). In Toll-like receptor or IL-1 signaling, Myd88-dependent NF-κB activation is linked by multiple signaling molecules, including TNF receptor associated factor 6 (TRAF6), inhibitor kappa B kinase α (IKKα), inhibitor kappa B kinase β (IKKβ), TGF-β-activated kinase 1 (TAK1), and TAK1-binding protein 2 (TAB2).^21^ We next used an NF-κB reporter gene to screen for key activation sites in inflammatory pathways. The results showed that ASB3 activated NF-κB reporter gene activity mediated only by TRAF6 but not by Myd88 and IKKβ (Figure 5b). We also found that overexpression of ASB3 upregulated IκBα phosphorylation mediated by TRAF6 (Figure 5c). TRAF6 is most widely described as a positive regulator of NF-κB signaling, and we speculate that ASB3 involvement in NF-κB activation may involve TRAF6 targeting. We then transfected labeled ASB3 and other signaling molecules into 293T cells and identified the proteins interacting with ASB3 by immunoprecipitation. TRAF6 protein, involved in the NF-κB signaling pathway, was identified as an interaction partner of ASB3 (Figure 5d). Furthermore, confocal imaging confirmed the specific colocalization of ASB3 and TRAF6 (Figure 5e). We also used *in vitro* transcription to further demonstrate that ASB3 interacts explicitly with TRAF6 but not TRAF3 (Figure 5f). The same phenomenon was also confirmed in colonic tissue (Figure S3a, b, Supporting Information). Previous studies have shown that TRAF6 contains an RF domain, a COIL domain, a ZnF domain, and a TRAF-C domain.^22^ ASB3 is composed of two main functional structure domains, including the ANK domain and the SOCX domain. We found that ASB3 interacts only with the C-terminal structural domain of TRAF6 (Figure 5g). Surprisingly, both the ANK domain and the SOCX domain were able to interact with TRAF6 (Figure 5h). Taken together, these results demonstrate that TRAF6 is essential for ASB3-mediated NF-κB activation.

**Figure. 5.**
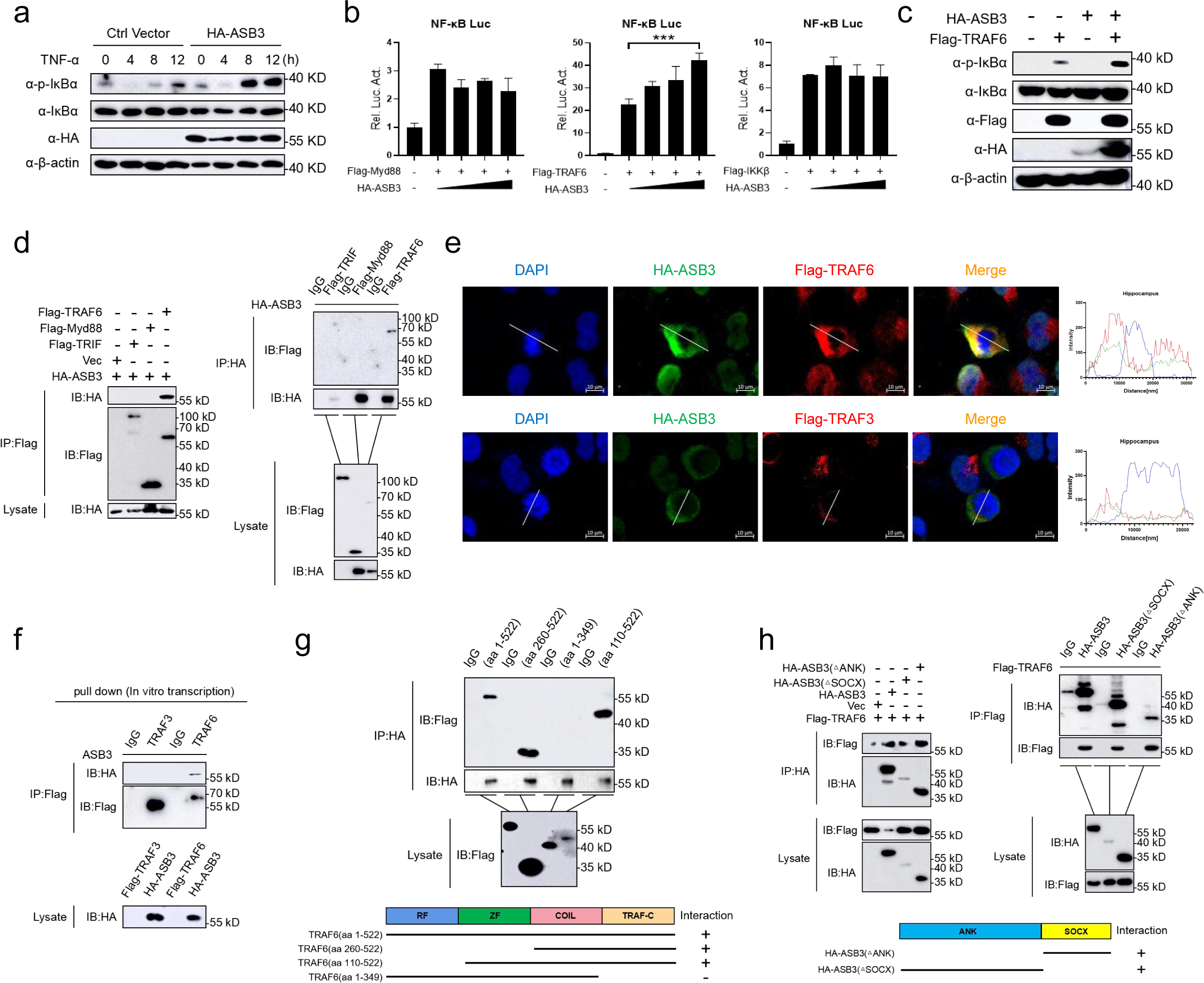
ASB3 specifically interacts with TRAF6. **a** HEK293T cells were transfected with HA-ASB3 plasmid or a vector control. 24 h after transfection, the cells were treated with TNF-α (150 ng/ml) for the indicated times. Cell lysates were separated by SDS-PAGE and analyzed by immunoblotting with the indicated antibodies. **b** HEK293T cells were transfected with the indicated plasmids along with control vector or increased amounts of ASB3 expression plasmids. Reporter assays were performed 24 h after transfection. **c** HEK293T cells were cotransfected with HA-ASB3 and Flag-TRAF6 plasmids or a vector control. **d, f-h** The indicated plasmids were transfected into HEK293T cells or transcribed *in vitro*. Then, coimmunoprecipitation and immunoblotting analyses were performed with the indicated antibodies. **e** HEK293T cells were transfected with HA-ASB3, Flag-TRAF6 or Flag-TRAF3 plasmids for 24 h and then stained with Flag antibody or HA primary antibody and secondary antibody. The nuclei were stained with DAPI. The fluorescence intensity profile of DAPI (blue), HA-ASB3 (green) and Flag-TRAF6 or Flag-TRAF3 (red) was measured along the line drawn by ZEN Blue. Scale bars, 50 µm. Difference were determined in **b** by unpaired Student’s t test.

### 2.6. ASB3 destabilizes TRAF6 by enhancing K48-linked polyubiquitination

We previously observed that overexpression of ASB3 specifically upregulated TRAF6-mediated NF-κB reporter gene activity. We hypothesized that ASB3 might regulate the stability of TRAF6. To test this assumption, we cotransfected HA-tagged ASB3 with flag-tagged Myd88, TRAF6, or IKKβ and performed immunoblotting analysis. Overexpression of ASB3 specifically downregulated TRAF6 expression but not TAB1 and IKKβ expression (Figure 6a). The ubiquitin-proteasome and autophagy-lysosome pathways are the major systems controlling protein degradation in eukaryotic cells.^23^ We found that ASB3-mediated TRAF6 degradation was restored only by the proteasome inhibitor MG132 but not by the autophagy inhibitor 3-methylamide (3-MA), the lysosomal inhibitor ammonium chloride (NH_4_Cl), or the pan-Caspase inhibitor ZVAD (Figure 6b).

**Figure. 6.**
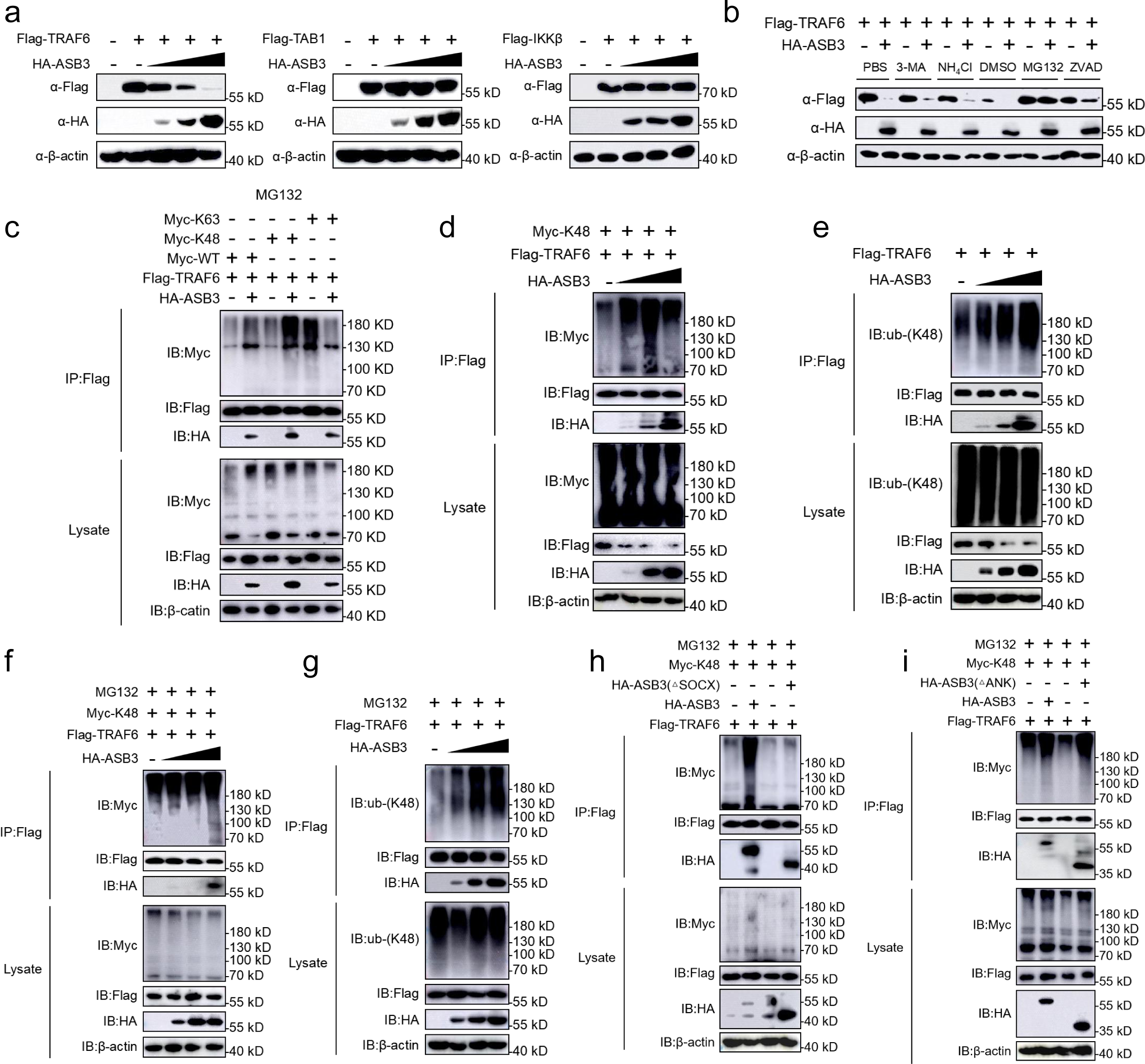
ASB3 promotes the K48-linked polyubiquitination of TRAF6. **a** HEK293T cells transfected with Flag-TRAF6, Flag-TAB1, or Flag-IKKβ with increasing amounts of HA-ASB3 for 24 h before immunoblotting analysis. **b** HEK293T cells were transfected with the indicated plasmids for 20 h and then treated with 3-methyladenine (3-MA) (10 mM), NH_4_Cl (20 mM), MG132 (10 μM), or ZVAD (20 μM) for 6 h. The cell lysates were then analyzed by immunoblotting with the indicated antibodies. **c** HEK293T cells transfected with Flag-TRAF6, Myc-ubiquitin, or its mutants [KO, which all but one lysine residues were simultaneously mutated to arginines (K-only)] together with control and ASB3 plasmids were pre-treated with MG132 (20 μM) for 6h. The cells were then subjected to coimmunoprecipitation and immunoblotting analysis with the indicated antibodies. **d** HEK293T cells transfected with Flag-TRAF6 and Myc-K48 with the increased amount of HA-ASB3 for 24 h. **e** HEK293T cells transfected with Flag-TRAF6 with the increased amount of HA-ASB3 for 24 h. **f** HEK293T cells transfected with Flag-TRAF6, Myc-K48 together with control and ASB3 plasmids were pretreated with MG132 (20 μM) for 6 h, and the cells were then subjected to coimmunoprecipitation and immunoblotting analysis with the indicated antibodies. **g** HEK293T cells transfected with Flag-TRAF6 with increasing amounts of HA-ASB3 for 20 h and then treated with MG132 (20 μM) for 6 h. **h, i** HEK293T cells were transfected with the indicated truncated plasmids for 20 h and then treated with or without MG132 (20 μM) for 6 h, and samples were collected at the indicated times. The cells were use in ubiquitination assays with the indicated antibodies.

To determine which ubiquitin chain of TRAF6 is modulated by ASB3, we analyzed ASB3-mediated ubiquitin mutants and the degradation of TRAF6. We found that ASB3 strongly mediates K48-linked polyubiquitination of TRAF6 in a mammalian overexpression system and in an IBD sample, but did not potentiate K63-linked ubiquitination of TRAF6 (Figure 6c-e) (Figure S4, Supporting Information). Adding MG132 led to the consistent conclusion that although TRAF6 protein expression was restored, the K48-linked polyubiquitin chains still accumulated (Figure 6f, g). To further explore which structural domain of ASB3 is required for enhanced K48-linked polyubiquitination of TRAF6, we continued to use ASB3 truncates and examined their effect on TRAF6 ubiquitination. The results showed that removing the SOCX domain eliminated ASB3-mediated endogenous K48-linked polyubiquitination of TRAF6 (Figure 6h). In contrast, truncation of the ANK domain does not eliminate TRAF6 exogenous K48-linked polyubiquitination (Figure 6i). In conclusion, ASB3 enhances K48-linked polyubiquitination of TRAF6 through the interaction of its SOCX domain with the TRAF6 C-terminal domain.

### 2.7. ASB3 disrupts TRAF6 stability in IECs exacerbates lethal inflammation

Since we observed that ASB3 deficiency eliminates K48-linked polyubiquitination of TRAF6 in mouse colon tissue (Figure 7a), we hypothesize that the mechanism of destabilizing regulation of TRAF6 by ASB3 may occur in IECs. Next, we constructed a model of colonic organoid inflammation and found that after TNF-α treatment, organoids from WT mice exhibited abnormal cell morphology and attenuated TRAF6 protein compared to those from ASB3^-/-^ mice (Figure S5a, Supporting Information) (Figure 7b). qPCR analysis showed that deletion of ASB3 in organoids was effective in alleviating the overexpression of proinflammatory factors (Figure 7c). To further confirm the effect of ASB3 on TRAF6 in IECs, we examined the types of ubiquitinated modifications. The results showed that ASB3 expression enhanced K48-linked polyubiquitination of TRAF6 in the organoid (Figure 7d). In addition, similar results were obtained with IECs dissociated from mouse colon tissue (Figure 7e). Collectively, these results reveal that ASB3 triggers massive release of proinflammatory factors from intestinal epithelial cells by targeting TRAF6 destabilization.

**Figure. 7.**
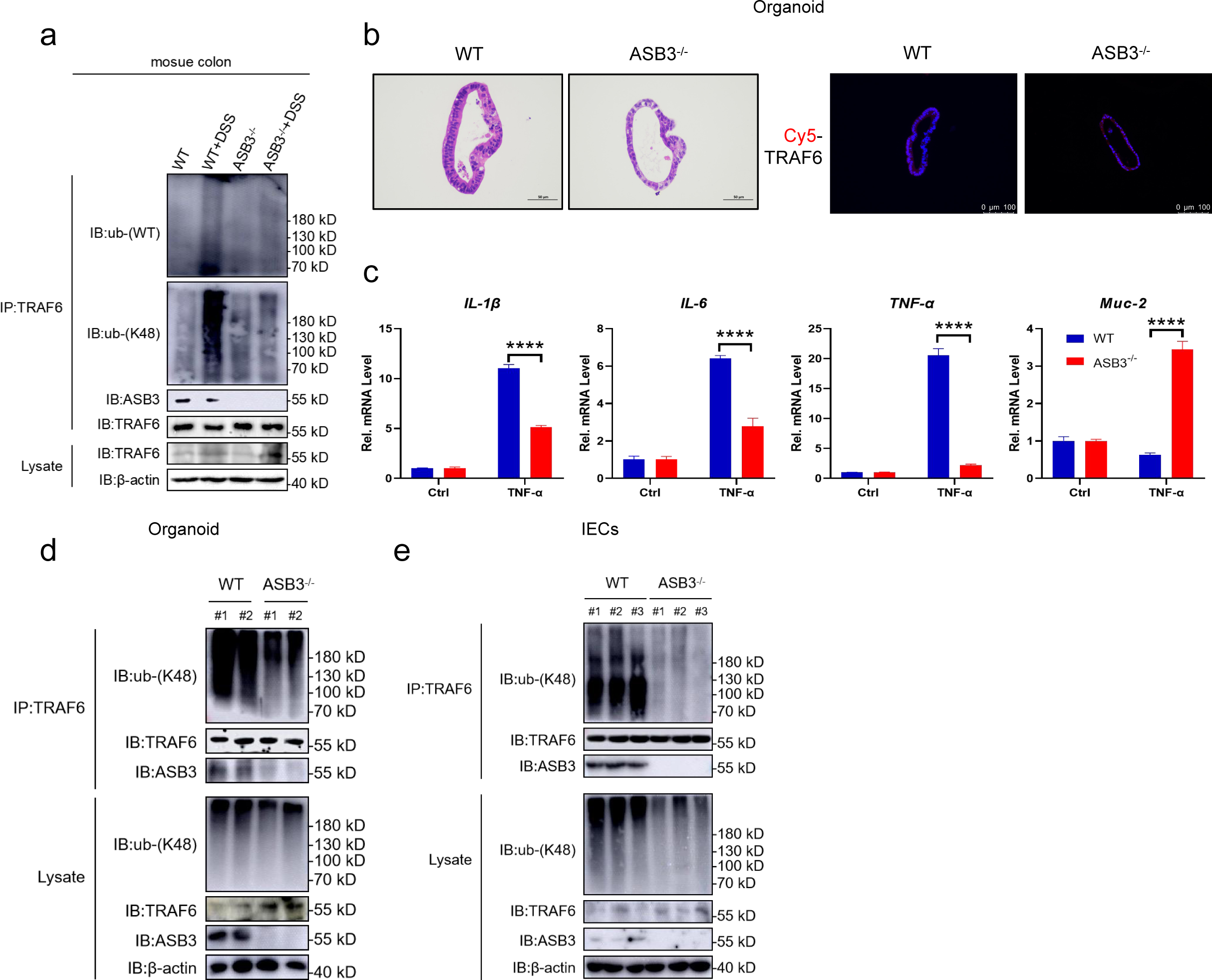
ASB3 promotes NF-κB signaling by limiting TRAF6 stability in IECs. **a** Total colonic tissue proteins from DSS-treated WT and ASB3^-/-^ mice. The samples were use in ubiquitination assays with the indicated antibodies. **b** Representative H&E staining and IF in the organoid. Scale bar, 50 μm or 100 μm. **c** The mRNA expression of IL-1β, IL-6, TNF-α, and Muc-2 was measured in both TNF-α-treated organoids relative to controls. **d** Total organoid proteins from TNF-α-treated WT and ASB3^-/-^ mice. **e** Total protein of IECs from dissociated colonic tissue from DSS-treated WT and ASB3^-/-^ mice. The samples were used in ubiquitination assays with the indicated antibodies. Difference were determined in **c** by One-way ANOVA.

## 3. Discussion

Ubiquitination plays an essential role in the pathogenesis and development of IBD. Ubiquitin-modifying enzymes (UMEs) coordinate the optimal ubiquitination of target proteins through synergistic effects, thereby maintaining gut homeostasis. However, in some reactions, these UMEs are aberrantly expressed and thus play a negative role. In the present study, we demonstrated that excessive activation of the E3 ligase ASB3 promotes the secretion of proinflammatory factors. In addition, we identified another key role of ASB3 in regulating gut microbiota imbalance by amplifying inflammatory signals. Mechanistically, ASB3 specifically enhanced K48-linked polyubiquitination, leading to TRAF6 destabilization. In this manner, proinflammatory factors are released in large quantities, the intestinal microecology and immune system are modulated, and damage due to inflammation is exacerbated. Thus, this study reveals a novel strategy for maintaining intestinal homeostasis by ASB3 protein-regulated inflammatory responses (Figure 8).

**Figure. 8.**
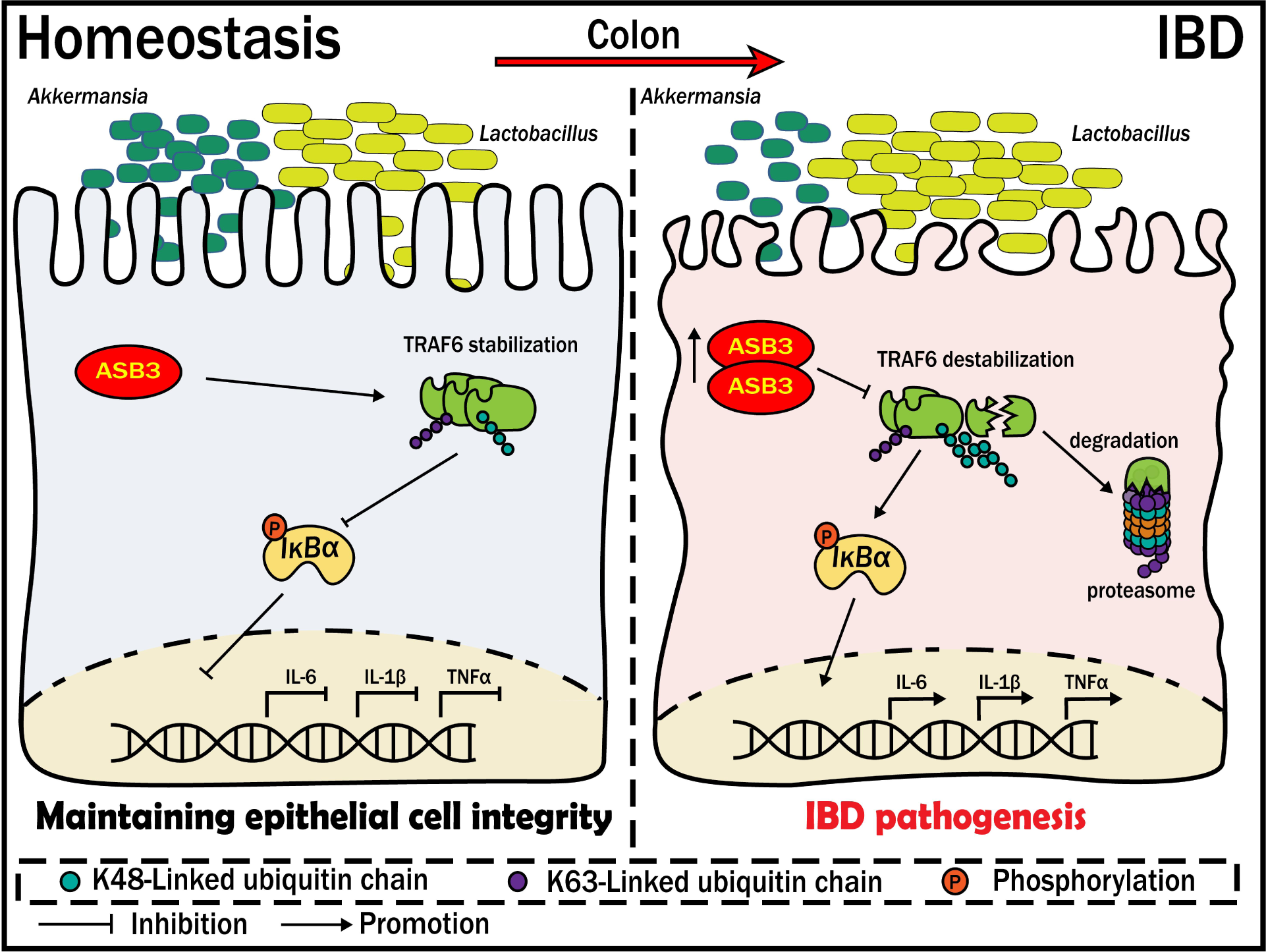
ASB3 aggravates intestinal epithelial inflammation and perturbs gut microbiota homeostasis by targeting TRAF6 destabilization. Under pathological conditions, ASB3 is upregulated in intestinal epithelial cells, leading to TRAF6 degradation via augmented K48-linked polyubiquitination. This results in the activation of TLR-independent NF-κB signaling, compromising the integrity of the intestinal epithelium and exacerbating the imbalance between the protective probiotic *Akkermansia* and *Lactobacillus*. Our study underscores the significance of ASB3 as a crucial mediator for preserving intestinal mucosal homeostasis by interacting with TRAF6.

A significant enhanced in ASB3 expression was reported in foci sites from UC and CD patients, compared to healthy sites (Figure 1a-d, j). This phenomenon of exacerbated mucosal inflammation was confirmed in both our animal and organoid models (Figure 1e-h). The role of ASB3 in intestinal diseases is poorly studied and is currently only found to be upregulated in COAD (Figure S1a, Supporting Information).^24^ However, one study reported that ASB3 could inhibit CRC metastasis by delaying the epithelial-mesenchymal transition.^25^ Our current studies showed that ASB3^-/-^ mice had downregulated susceptibility to experimental colitis and preserved intestinal barrier function (Figure 2). Interestingly, enhanced ASB3 expression is causally related to impaired intestinal homeostasis by the massive secretion of colonic proinflammatory factors. Thus, aberrant expression of ASB3 may have significant pathophysiological consequences.

ASB3 overexpression inhibited TRAF6 protein stability (Figure 6a) but not transcription (Figure S4a, Supporting Information). We found that ASB3 specifically interacted with TRAF6 and degraded TRAF6 through K48-linked polyubiquitination (Figure 6c-g). A previous study has reported that increased expression of zinc finger protein A20 inhibited TRAF6/NF-κB activation.^26^ TRAF6 autophagic degradation has also been shown to block NF-κB signaling.^27^ This may seem paradoxical at first; TRAF6 destabilization in our study still resulted in NF-κB expression and inflammatory factor secretion. Surprisingly, however, we noted in one experiment that mice lacking TRAF6 in IECs exhibited exacerbated DSS-induced inflammatory responses that TLR-related signaling molecules (Myd88 and TRIF) did not trigger.^28^ TRAF6 signaling independent of TLRs likely limits DSS-induced colitis not by mediating intrinsic functions involved in IECs homeostasis, but instead by inducing the release of epithelial cytokines such as TGF-β to exert direct or indirect anti-inflammatory functions.^29^ MALT1 has been reported to regulate STAT3 during IBD recovery to positively promote mucosal healing.^30^ This explains the upregulation of IL-22 expression in the colon, which may be achieved through interactions between IECs and ILCs. Numerous studies have shown that the TLR-induced NF-κB signaling pathway depends on regulation of E3 ligases.^31^ A20 was shown to be an important regulator of NF-κB and to have a bidirectional function in inflammatory stabilization.^32^ In fact, we found that ASB3-mediated destabilization of TRAF6 activated NF-κB *in vivo* and *in vitro*, which may be a novel mechanism for disrupting the homeostasis of inflammation maintained by TRAF6.

In the present study, we found a negative correlation between ASB3 expression and Lgr5 (Figure S5b, Supporting Information). Recent reports have confirmed that the E3 ubiquitin ligase TRIM27 maintains intestinal homeostasis by activating Wnt signaling to promote the self-renewal of Lgr5^+^ ISCs.^33^ Intestinal homeostasis requires the integrity of the intestinal epithelial barrier to be maintained, and epithelial cells maintain dynamic homeostasis by coordinating with other intestinal cell populations and with cytokines.^34^ Whether ASB3 affects ISCs to drive epithelial cell proliferation and repair needs to be investigated in depth in the future with the help of organoid models. Immune cells in ASB3-deficient mice behaved normally at steady state, and intestinal ILC3 numbers were upregulated in inflammatory states (Figure S6, Supporting Information). Our study also showed more severe intestinal inflammation accompanied by a reduced amount of ILC3s, possibly limited by the degree of epithelial cell inflammation (Figure 3d). The immune-related roles played by intestinal epithelial cells are primarily driven by gut-resident natural immune cells. These cell populations depend on the regulation of several molecules during IBD. DR3-driven loss of ILC3s in the large intestine is an essential factor contributing to the exacerbation of colitis,^35^ and mice lacking CD93 expression in ILC3s also exhibit impaired IL-22 production and increased colonic inflammation.^36^ In another study, metabolic reprogramming during intestinal LTi cell or ILC3 activation was dependent on the regulation of the immune checkpoint PD-1.^37^ ASB3 has been reported to degrade TNF-R2 via the proteasome pathway,^8^ and whether there is a correlation between this and DR3 remains to be explored. Our data confirm that targeting ASB3/TRAF6 interactions largely affects the tissue repair function of ILC3s by disrupting the intestinal barrier. However, we currently have no direct evidence to suggest that ASB3 is involved in ILC3 regulation.

We observed that ASB3-deficient mice acquired colitis resistance by protecting more Bacteroidetes species while reducing the sudden increase in *Lactobacillus* (Figure S2, Supporting Information). Intestinal immunity and homeostasis are regulated by epithelial cell/immune cell interactions and require fine-tuned communication between the host and microbes. The growth of certain pathogenic bacteria usually manifests as IBD-associated microbial dysbiosis at the cost of loss of protective commensal bacteria. Host genetic mutations often amplify this disorder. ANG1 deficiency antagonizes the growth of Lachnospiraceae by increasing the level of α*-Proteobacteria*, leading to dysbiosis of the intestinal ecology.^38^ Similarly, Nlrp12 deficiency led to a reduction in microbiota diversity, loss of protective intestinal commensal strains of Lachnospiraceae, and an increase in the highly pathogenic bacterium Erysipelotrichaceae.^5^ CARD9 deletion reduces members of *L. reuteri*, *Allobaculum*, and the phylum Actinobacteria and weakens intestinal tryptophan catabolic function.^39^ Although the gut microbiota changes underlying IBD are not unique, most of the Bacteroidetes species identified in one study were depleted in IBD, and the abundance of *Streptococcus* and *Lactobacillus* was increased.^40^ Members of Muribaculaceae and Lachnospiraceae have been identified as significant utilizers of mucin monosaccharides, and members of Rikenomycetaceae and Bacteroidetaceae are also thought to have functions in the utilization of mucin monosaccharides.^41^ Interconversion of mucin monosaccharide-utilizing intestinal commensal bacteria caused by ASB3 deficiency before and after DSS challenge also implies that ASB3 may have potential effects on the intestinal mucus layer.

In this study we report a previously unidentified function of ASB3 deficiency in ameliorating DSS-induced colitis. Our data suggest that high expression of ASB3 in intestinal epithelial cells under pathological conditions mediates TRAF6 destabilization, thereby inducing proinflammatory factor release and impaired intestinal barrier. ASB3 exacerbates the interaction between abnormal immune signaling and dysbiotic microbiota, creating a vicious cycle that leads to persistent inflammation and ecological dysregulation in the gut. This evidence increases our understanding of the pathogenesis of colitis. Interventions of ASB3-mediated proinflammatory pathway and ASB3-synergistic microbiota may have potential relevance for clinical IBD applications.

## 4. Experimental Section

### Human Samples

Human colon samples from IBD patients were provided by the tissue bank in accordance with its regulations and approved by the Ethics Committee of Tongji University and Shanghai Tenth People’s Hospital.

### Mice

ASB3^-/-^ mice were generated using PiggyBac transposon-based targeting technology and were kindly provided by Professor Dongqin Yang (Shanghai Fudan University). Mice used in this study were housed under specific-pathogen-free (SPF) conditions and had an FVB genetic background. To avoid the effect of familial heritability on the colony, ASB3^-/-^ mice and the wild-type (WT) controls were all generated from the same heterozygous (HE) ASB3^+/-^ parents. WT, HE, and ASB3^-/-^ mice were identified by PCR amplification and DNA sequencing with the primers 5′-CTGAGATGTCCTAAATGCACAGCG-3′, 5′-CCATGACCAAACCCAATTTACACAC-3′, and 5′-TTGTTTTTTTTCCCCCCTAGACAGG-3′, respectively. For the cohousing experiment, 4-week-old mice from the same mother were divided into either individually reared (SiHo) or cohoused with age- and sex-matched mice (CoHo) for 6 weeks. For antibiotic treatment, mice were given an ABX cocktail (33.2 mg ampicillin, 33.2 mg neomycin, 33.2 mg metronidazole, and 16.7 mg vancomycin) orally for 5 consecutive days.^42^

### Induction of DSS-mediated acute colitis

Six- to eight-week-old sex-matched mice were induced to develop acute colitis with 3.0% (w/v) DSS (MW 36-50 kDa; MP Biomedical) dissolved in sterile, distilled water ad libitum for experimental days 1 to 6 and were then provided regular water for another 2 days.^43^ The DSS solution was made freshly every day.

### Cells, organoids and plasmids

Human embryonic kidney cells (HEK293T) and human colorectal adenocarcinoma cells (HT-29) were cultured in Dulbecco’s modified Eagle’s medium (DMEM) and RPMI-1640 supplemented with 10% (v/v) fetal bovine serum (FBS) and 2 mM glutamine with penicillin (100 U/ml)/streptomycin (100 mg/ml). Intestinal epithelial cells (IECs) or crypts were dissociated from colonic segments of WT or ASB3^-/-^ mice. Isolated colonic crypts were resuspended in DMEM/F12 medium, counted and resuspended in colonic organoid growth medium and matrix gel (Corning) at a 1:1 ratio.^44^ The medium was changed every other day, and after 7 days, the organoid was stimulated with 150 ng/ml mouse recombinant TNF-α (PeproTech) for 24 h.^45^ All cells were cultured and maintained at 37 °C with 5% CO_2_.

Plasmids for Flag-tagged Myd88, TRIF, TRAF3, TRAF6, TAB1, IKKβ, Myc-tagged ubiquitin (Myc-K48), NF-κB-Luc and pRL-TK (internal control luciferase reporter plasmid) used in the study were described previously.^23^ Plasmids for HA-tagged ASB3, ASB3 (ΔANK), and ASB3 (ΔSOCX) were kindly provided by Prof. Dongqin Yang (Fudan University, Shanghai, China). Plasmids for Flag-tagged TRAF6 (aa 260-522), TRAF6 (aa 110-522), and TRAF6 (aa 1-349) were kindly provided by Prof. Qiyun Zhu (Lanzhou Veterinary Research Institute, Chinese Academy of Agricultural Sciences, China).

### Histology

For histopathological analysis, colon tissue from WT and ASB3^-/-^ mice was fixed in 4% paraformaldehyde solution overnight for processing and embedded in paraffin wax according to standard procedures. Then, 3 μm sections were taken and stained with hematoxylin and eosin or Alcian blue. The stained sections were scanned with a microscope (Leica).

### Confocal microscopy

Confocal microscopy was performed as previously described.^46^ The indicated plasmids were cotransfected into 293T cells and collected after 24 h. Cells were fixed with 4% paraformaldehyde for 15 min at room temperature and subsequently blocked and permeabilized with 0.5% Triton X-100 5% skim milk 1 h at 4 °C. After blocking, slides for immunofluorescence were incubated with fluorescein-conjugated secondary antibodies (1:500) overnight at 4°C in the dark and mounted with DAPI (Beyotime) (1:1000). Images were acquired with a Zeiss microscope (LSM 710) to visualize stained cells.

### Statistical analysis

Data are presented as the mean ± SD unless otherwise indicated. All samples were analyzed using GraphPad Prism 8 software. Data were analyzed by unpaired two-tailed Student’s t test or two-way ANOVA with Sidak’s test to correct for multiple comparisons. Probability (p) values of < 0.05 were considered significant: \**P* < 0.05, \*\**P* < 0.01, \*\*\**P* < 0.001 and \*\*\*\**P* < 0.0001; n.s., not significant.

## Data availability

The 16S rRNA sequencing of the fecal microbiota has been deposited on the Sequence Read Archive website with the BioProject accession number PRJNA1000707 (Temporary Submission ID: SUB13720433).

## Acknowledgments

We thank Dr. Bin Xu (Department of General Surgery, Shanghai 10th People’s Hospital, Tongji University) for donating the CD patients sample. We thank Prof. Dongqin Yang (Central laboratory, Huashan Hospital, Fudan University) for donating the UC patients samples, ASB3^+/-^ mice and HA-tagged ASB3 plasmids. We thank Prof. Qiyun Zhu (Lanzhou Veterinary Research Institute, Chinese Academy of Agricultural Sciences) for donating the Flag-tagged TRAF6 truncated plasmids.

## Author contributions

MC performed the majority of the experiments. BX, YS, JW, YL, CS, WZ, XL and XS contributed and/or supported us for some experiments. BX, and DY provided/analyzed human material/samples. XC, YZ and MC contributed toward the animal studies and wrote the manuscript. JW, NW, WY, YJ and HH participated in the inflammation evaluation experiments. MC and BX analysed and interpreted data. MC, YZ, GY, CW and XC conceived and designed the experiments, interpreted the data and wrote the manuscript. XC acts as guarantor for the present study. All authors discussed the results and approved the final manuscript.

## References

1 Shan, Y., Lee, M. & Chang, E. The Gut Microbiome and Inflammatory Bowel Diseases. Annual review of medicine 73, 455–468, doi:10.1146/annurev-med-042320-021020 (2022).

2 Mehandru, S. & Colombel, J. The intestinal barrier, an arbitrator turned provocateur in IBD. Nature reviews. Gastroenterology & hepatology 18, 83–84, doi:10.1038/s41575-020-00399-w (2021).

3 Caruso, R., Lo, B. & Núñez, G. Host-microbiota interactions in inflammatory bowel disease. Nature reviews. Immunology 20, 411–426, doi:10.1038/s41577-019-0268-7 (2020).

4 Glassner, K., Abraham, B. & Quigley, E. The microbiome and inflammatory bowel disease. The Journal of allergy and clinical immunology 145, 16–27, doi:10.1016/j.jaci.2019.11.003 (2020).

5 Chen, L. et al. NLRP12 attenuates colon inflammation by maintaining colonic microbial diversity and promoting protective commensal bacterial growth. Nature immunology 18, 541–551, doi:10.1038/ni.3690 (2017).

6 Zhang, W. et al. ASB3 knockdown promotes mitochondrial apoptosis via activating the interdependent cleavage of Beclin1 and caspase-8 in hepatocellular carcinoma. Science China. Life sciences 62, 1692–1702, doi:10.1007/s11427-018-9505-0 (2019).

7 Li, R. et al. E3 ligase ASB8 promotes porcine reproductive and respiratory syndrome virus proliferation by stabilizing the viral Nsp1α protein and degrading host IKKβ kinase. Virology 532, 55–68, doi:10.1016/j.virol.2019.04.004 (2019).

8 Chung, A., Guan, Y., Yuan, Z., Albina, J. & Chin, Y. Ankyrin repeat and SOCS box 3 (ASB3) mediates ubiquitination and degradation of tumor necrosis factor receptor II. Molecular and cellular biology 25, 4716–4726, doi:10.1128/mcb.25.11.4716-4726.2005 (2005).

9 Ruan, J., Schlüter, D., Naumann, M., Waisman, A. & Wang, X. Ubiquitin-modifying enzymes as regulators of colitis. Trends in molecular medicine 28, 304–318, doi:10.1016/j.molmed.2022.01.006 (2022).

10 Xiao, Y., Huang, Q., Wu, Z. & Chen, W. Roles of protein ubiquitination in inflammatory bowel disease. Immunobiology 225, 152026, doi:10.1016/j.imbio.2020.152026 (2020).

11 Hou, P. et al. An unconventional role of an ASB family protein in NF-κB activation and inflammatory response during microbial infection and colitis. Proceedings of the National Academy of Sciences of the United States of America 118, doi:10.1073/pnas.2015416118 (2021).

12 Lin, H. et al. The membrane-associated E3 ubiquitin ligase MARCH3 downregulates the IL-6 receptor and suppresses colitis-associated carcinogenesis. Cellular & molecular immunology 18, 2648–2659, doi:10.1038/s41423-021-00799-1 (2021).

13 Lian, Q. et al. TRIM34 attenuates colon inflammation and tumorigenesis by sustaining barrier integrity. Cellular & molecular immunology 18, 350–362, doi:10.1038/s41423-020-0366-2 (2021).

14 Yao, Y. et al. Mucus sialylation determines intestinal host-commensal homeostasis. Cell 185, 1172–1188.e1128, doi:10.1016/j.cell.2022.02.013 (2022).

15 Sayed, I. et al. Host engulfment pathway controls inflammation in inflammatory bowel disease. The FEBS journal 287, 3967–3988, doi:10.1111/febs.15236 (2020).

16 Fan, W. et al. Estrogen receptor β activation inhibits colitis by promoting NLRP6-mediated autophagy. Cell reports 41, 111454, doi:10.1016/j.celrep.2022.111454 (2022).

17 Diefenbach, A., Gnafakis, S. & Shomrat, O. Innate Lymphoid Cell-Epithelial Cell Modules Sustain Intestinal Homeostasis. Immunity 52, 452–463, doi:10.1016/j.immuni.2020.02.016 (2020).

18 Wang, B. et al. Macrophage β2-Integrins Regulate IL-22 by ILC3s and Protect from Lethal Citrobacter rodentium-Induced Colitis. Cell reports 26, 1614–1626.e1615, doi:10.1016/j.celrep.2019.01.054 (2019).

19 Zhao, Q. & Maynard, C. Mucus, commensals, and the immune system. Gut microbes 14, 2041342, doi:10.1080/19490976.2022.2041342 (2022).

20 Wan, P. et al. ASB17 Facilitates the Burst of LPS-Induced Inflammation Through Maintaining TRAF6 Stability. Frontiers in cellular and infection microbiology 12, 759077, doi:10.3389/fcimb.2022.759077 (2022).

21 Hayden, M. & Ghosh, S. Shared principles in NF-kappaB signaling. Cell 132, 344–362, doi:10.1016/j.cell.2008.01.020 (2008).

22 Chen, Z. et al. Influenza D virus Matrix protein 1 restricts the type I interferon response by degrading TRAF6. Virology 568, 1–11, doi:10.1016/j.virol.2022.01.001 (2022).

23 Zeng, Y. et al. The PB1 protein of influenza A virus inhibits the innate immune response by targeting MAVS for NBR1-mediated selective autophagic degradation. PLoS pathogens 17, e1009300, doi:10.1371/journal.ppat.1009300 (2021).

24 Mu, L. et al. Pan-cancer analysis of ASB3 and the potential clinical implications for immune microenvironment of glioblastoma multiforme. Frontiers in immunology 13, 842524, doi:10.3389/fimmu.2022.842524 (2022).

25 Du, W. et al. The loss-of-function mutations and down-regulated expression of ASB3 gene promote the growth and metastasis of colorectal cancer cells. Chinese journal of cancer 36, 11, doi:10.1186/s40880-017-0180-0 (2017).

26 Deng, H. et al. A20 Establishes Negative Feedback With TRAF6/NF-κB and Attenuates Early Brain Injury After Experimental Subarachnoid Hemorrhage. Frontiers in immunology 12, 623256, doi:10.3389/fimmu.2021.623256 (2021).

27 Deng, T. et al. avibirnavirusTRAF6 autophagic degradation by VP3 inhibits antiviral innate immunity via blocking NFKB/NF-κB activation. Autophagy 18, 2781–2798, doi:10.1080/15548627.2022.2047384 (2022).

28 Vlantis, K. et al. TLR-independent anti-inflammatory function of intestinal epithelial TRAF6 signalling prevents DSS-induced colitis in mice. Gut 65, 935–943, doi:10.1136/gutjnl-2014-308323 (2016).

29 Beck, P. et al. Transforming growth factor-beta mediates intestinal healing and susceptibility to injury in vitro and in vivo through epithelial cells. The American journal of pathology 162, 597–608, doi:10.1016/s0002-9440(10)63853-9 (2003).

30 Wittner, L. et al. Proteolytic Activity of the Paracaspase MALT1 Is Involved in Epithelial Restitution and Mucosal Healing. International Journal of Molecular Sciences 24, 7402 (2023).

31 Lu, Y. et al. RING finger 138 deregulation distorts NF-кB signaling and facilities colitis switch to aggressive malignancy. Signal transduction and targeted therapy 7, 185, doi:10.1038/s41392-022-00985-1 (2022).

32 Yu, X. et al. MYD88 L265P elicits mutation-specific ubiquitination to drive NF-κB activation and lymphomagenesis. Blood 137, 1615–1627, doi:10.1182/blood.2020004918 (2021).

33 Wang, J. et al. TRIM27 maintains gut homeostasis by promoting intestinal stem cell self-renewal. Cellular & molecular immunology 20, 158–174, doi:10.1038/s41423-022-00963-1 (2023).

34 Xie, Y. et al. Gut epithelial TSC1/mTOR controls RIPK3-dependent necroptosis in intestinal inflammation and cancer. The Journal of clinical investigation 130, 2111–2128, doi:10.1172/jci133264 (2020).

35 Li, J. et al. Activation of DR3 signaling causes loss of ILC3s and exacerbates intestinal inflammation. Nature communications 10, 3371, doi:10.1038/s41467-019-11304-8 (2019).

36 Huang, J. et al. Interleukin-17D regulates group 3 innate lymphoid cell function through its receptor CD93. Immunity 54, 673–686.e674, doi:10.1016/j.immuni.2021.03.018 (2021).

37 Wu, D. et al. PD-1 signaling facilitates activation of lymphoid tissue inducer cells by restraining fatty acid oxidation. Nature metabolism 4, 867–882, doi:10.1038/s42255-022-00595-9 (2022).

38 Sun, D. et al. Angiogenin maintains gut microbe homeostasis by balancing α-Proteobacteria and Lachnospiraceae. Gut 70, 666–676, doi:10.1136/gutjnl-2019-320135 (2021).

39 Lamas, B. et al. CARD9 impacts colitis by altering gut microbiota metabolism of tryptophan into aryl hydrocarbon receptor ligands. Nature medicine 22, 598–605, doi:10.1038/nm.4102 (2016).

40 Schirmer, M., Garner, A., Vlamakis, H. & Xavier, R. Microbial genes and pathways in inflammatory bowel disease. Nature reviews. Microbiology 17, 497–511, doi:10.1038/s41579-019-0213-6 (2019).

41 Pereira, F. et al. Rational design of a microbial consortium of mucosal sugar utilizers reduces Clostridiodes difficile colonization. Nature communications 11, 5104, doi:10.1038/s41467-020-18928-1 (2020).

42 Li, H., et al. Bifidobacterium spp. and their metabolite lactate protect against acute pancreatitis via inhibition of pancreatic and systemic inflammatory responses. Gut microbes 14, 2127456, doi:10.1080/19490976.2022.2127456 (2022).

43 Cui, S. et al. CD1d1 intrinsic signaling in macrophages controls NLRP3 inflammasome expression during inflammation. Science advances 6, doi:10.1126/sciadv.aaz7290 (2020).

44 Miyoshi, H. & Stappenbeck, T. In vitro expansion and genetic modification of gastrointestinal stem cells in spheroid culture. Nature protocols 8, 2471–2482, doi:10.1038/nprot.2013.153 (2013).

45 Zhang, J. et al. MPST deficiency promotes intestinal epithelial cell apoptosis and aggravates inflammatory bowel disease via AKT. Redox biology 56, 102469, doi:10.1016/j.redox.2022.102469 (2022).

46 Cheng, M. et al. African Swine Fever Virus L83L Negatively Regulates the cGAS-STING-Mediated IFN-I Pathway by Recruiting Tollip To Promote STING Autophagic Degradation. Journal of virology 97, e0192322, doi:10.1128/jvi.01923-22 (2023).

