## Supplementary Methods for "ASB3 expression aggravates inflammatory bowel disease by targeting TRAF6 protein stability and affecting the intestinal microbiota"

*Induction of DSS-mediated acute colitis*

6- to 8-week-old sex-matched mice were induced to develop acute colitis by 3.0% (w/v) DSS (MW 36-50 kDa; MP Biomedical) dissolved in sterile, distilled water ad lib for the experimental days 1 to 6, and were then provided regular water for another 2 days.(1) The DSS solution was made freshly every days. The severity of colitis was assessed by the Disease Activity Index (DAI), including daily weight loss, stool consistency, and occult blood, as previously described.(2) Briefly, weight score is as follows: 0 means no weight loss, 1 means 1-5% weight loss, 2 means 5-10% weight loss 3 means 10-20% weight loss, 4 means more than 20% weight loss. The consistency of the stool is scored as follows: 0 indicates normal stool pattern, 1 indicates semi-formed stool not adhering to the anus, 2 indicates semi-formed stool adhering to the anus, and 3 indicates liquid stool adhering to the anus. Bleeding was scored as follows: 0 indicates blood occult negative, 1 indicates blood occult positive, 2 indicates blood traces in the stool; 3 indicates obvious blood traces in the anus, and 4 indicates rectal haemorrhage. Weight score, stool consistency score and bleeding score were added and expressed as clinical scores. The entire colon was removed on day 8 to measure the colon length.

*Antibodies and reagents*

The antibodies used in this study were as follows: HRP-conjugated anti-HA (12013819001), Myc (11814150001) antibodies (Roche); phosphorylated IκBα (AF5851), IκBα (AG2737), HRP-labeled Goat anti-Mouse IgG(H+L) (A0216), and HRP-labeled Goat anti-Rabbit IgG(H+L) (A0208) antibodies (Beyotime), FITC Goat Anti-Mouse IgG (H+L) (K1201), Cy5 Goat Anti-Rabbit IgG (H+L) (K1212) secondary antibodies (APExBIO); anti-TRAF3 (66310-1-Ig), anti-HA (66006-2-Ig), anti-DYKDDDDK (20543-1-AP), anti-β-actin (66009-1-Ig), and anti-Villin (16488-1-AP) antibodies (Proteintech); anti-TRAF6 antibody (8028), anti-ubiquitin (K48) (8081) and HRP-conjugated mouse anti-rabbit IgG (Conformation Specific) (5127) antibodies (Cell Signaling Technology); anti-ubiquitin (WT) antibody (Santa Cruz, sc-8017); anti-ASB3 (88812) antibody (MBL); anti-TRAF6 antibody (ab137452), anti-EpCAM antibody (ab71916) (Abcam); HRP-conjugated anti-Flag (A8592) antibody (Sigma). Reagents used in the study included: 3-Methyladenine (M9281), MG132 (M7449), DMSO (D2650), NH_4_Cl (09718), anti-Myc agarose affinity beads (A7470) and protein A/G agarose affinity beads (P6486/E3403) (Sigma); mouse IL-1β/IL-6/TNF-α ELISA kit (MM-0040M1/MM-0163M1/MM-0132M1) (MEIMIAN); mouse recombinant TNF-α (315-01A) (PeproTech); DAPI (C1005), ZVAD (Caspase inhibitor Z-VAD-FMK, C1202), NP-40 (ST366), HEPES Solution (C0215) (Beyotime). TnT® T7/SP6 Quick Coupled Transcription/Translation System (L1170/L2080) (Promega).

*Coimmunoprecipitation (Co-IP)*

HEK293T cells at 80-90% confluency were washed with PBS three times and collected using a cell scraper into tubes. The same mass of colonic tissue was washed three times with precooled PBS and mechanically homogenized with RIPA lysis buffer (Thermo Scientific) containing 1% PMSF. All samples were lysed in NP-40 lysis buffer containing 20 mM Tris-HCl (pH 8.0), 1 mM EDTA, 1% NP-40, and 150 mM NaCl supplemented with Halt Protease Inhibitor Cocktail and kept on ice for 30 min. After 30 min of incubation, the lysates were spun down at 12000 rpm at 4 °C for 20 min. All supernatants were collected, and protein was quantified using the BCA Protein Assay Kit (Beyotime). The supernatants were pretreated with 30 µl of anti-Flag agarose affinity gels or protein A/G at 4 °C for 2 h. The indicated primary antibody or IgG control (mouse IgG, Beyotime) was then added to the pretreated lysates and incubated overnight at 4 °C. The IP complexes and whole-cell lysates were subjected to protein transfer by the conventional Western blotting method, and protein development was detected using the above mentioned antibodies.

*cLPLs isolation and flow cytometry analysis*

The LPLs were isolated as described previously.(3) Briefly, the colon is opened longitudinally in pre-chilled PBS to expose the lumen of the tube. colon tissue was gently rinsed, the intestinal contents removed and cut into small 1 cm pieces. Tissue pieces were incubated in pre-digested medium (RPMI- 1640 containing 20 mM HEPES, 5 mM EDTA, 1% penicillin and streptomycin, 2% FBS, 2 mM DTT) for 20 min at 37 °C and 180 rpm in a shaking incubator. Shake vigorously for 2 min and wash with PBS. The remaining tissues were transferred to digestion buffer (RPMI-1640 supplemented with 1% penicillin and streptomycin, 20 mM HEPES, 1.5 mg/mL collagenase IV, 0.5 mg/mL DNase I and 2% FBS) and incubated in a shaking incubator at 37 °C, 250 rpm for two consecutive 15 min. Cells were washed and collected through a 70 µm cell membrane. After filtering, leukocytes were purified on Percoll density-gradient separation. Purified cLPs were washed and stimulated with 50 ng/mL PMA (Sigma) and 500 ng/mL ionomycin (Sigma) in the presence of GolgiPlug (BD) for 4 h prior to staining for flow cytometry analysis by LSR Fortessa (BD). Flurochrome-conjugated antibodies used in this study were as follows: Fixable Viability Stain 780 (565388), Biotin-labeled TCR β Chain (553168), Biotin-labeled γδT-Cell Receptor (553176), Biotin-labeled CD11b (557395), Biotin-labeled CD19 (553784), Biotin-labeled Ly-6G and Ly-6C (553124), Biotin-labeled TER-119 (553672), Streptavidin APC-Cy™7 (554063), CD16/32 (553140), CD45.1 (550994), CD3 (557984), CD4 (561115), CD8 (553035), CD44 (560567), CD25 (553075), Nkp46 (560756), CCR6 (557976), RORgt (562607) (BD Biosciences); IL-22 (516409) (Biolegend). Cell surface staining was performed by incubating cells with antibodies for 30 min at 4°C after blocking with CD16/32. RORgt and IL-22 staining was carried out using intracellular transcription factor kit (eBioscience) or cytokine staining kit (BD Biosciences).

*16S rDNA sequencing*

16S rDNA amplicon libraries were produced from DNA of Feces and was completed by Tianjin Novogene Bioinformatics Technology Co.Ltd. (Tianjin, China). Total genome DNA from samples was extracted using CTAB/SDS method. DNA concentration and purity was monitored on 1% agarose gels. According to the concentration, DNA was diluted to 1ng/µL using sterile water. PCR amplification of the V3-V4 region of the bacterial 16S rRNA gene was performed using specific primers with Barcode for the 16S V4 region primer 515F-806R based on the selection of the sequenced region. Sequencing libraries were generated using Illumina TruSeq DNA PCR-Free Library Preparation Kit (Illumina, USA) following manufacturer’s recommendations and index codes were added. The library quality was assessed on the Qubit@ 2.0 Fluorometer (Thermo Scientific) and Agilent Bioanalyzer 2100 system. At last, the library was sequenced on an Illumina NovaSeq platform and 250 bp paired-end reads were generated. The 16S rRNA sequencing of the fecal microbiota has been deposited on the Sequence Read Archive website with the BioProject accession number PRJNA1000707 (Temporary Submission ID: SUB13720433).

*Dual-luciferase reporter assays*

Plasmids encoding Flag-tagged MyD88, TRAF6, and IKKβ or HA-tagged ASB3, were co-transfected with NF-κB-Luc and pRL-TK into HEK293T cells for 24 h using Lipofectamine 3000 (Invitrogen) reagent. Cell samples were collected at the indicated times and lysed or assayed using the Dual-Luciferase® Reporter Gene (DLR™) Assay System (Promega) according to the manufacturer's instructions. Finally, the relevant reporter gene activity is assayed with Firefly luciferase and Renilla luciferase reagents.

*Quantitative real-time RT-PCR*

RNA was isolated with Trizol reagent (Takara). cDNA was synthesized using the reverse transcription Moloney mouse leukemia virus (M-MLV) reverse transcriptase (Promega). Real-time PCR was performed using SYBR Green Mix (Takara). Reactions were run with the Applied Biosystems 7500 real-time PCR System. The results were displayed as relative expression values normalized to GAPDH. Sequences of PCR primers are as follows: 5’-CTGATAACAGGGGATGGATCC (forward primer for Hu ASB3), 5’-CGAGATGCAAAGCACAGAAAC (reverse primer for Hu ASB3), 5’-ACAGAGGCTTACTCAGACACG (forward primer for mus ASB3), 5’-TCCCCTGTTATCAGCGACATC (reverse primer for mus ASB3), 5’-GCAACTGTTCCTGAACTCAACT (forward primer for mus IL-1b), 5’-ATCTTTTGGGGTCCGTCAACT (reverse primer for mus IL-1b), 5’-TCTGCAAGAGACTTCCATCCAGTTGC (forward primer for mus IL-6), 5’-AGCCTCCGACTTGTGAAGTGGT (reverse primer for mus IL-6), 5’-TCAGGTGCAAGGTGAAGTTG (forward primer for mus Reg3g), 5’-GGCCACTGTTACCACTGCTT (reverse primer for mus Reg3g), 5’-ATGCCCACCTCCTCAAAGAC (forward primer for mus Muc2), 5’-GTAGTTTCCGTTGGAACAGTGAA (reverse primer for mus Muc2), 5’-AAAATCAAGTGGGGCGATGCT (forward primer for Hu GAPDH), 5’-GGGCAGAGATGATGACCCTTT (reverse primer for Hu GAPDH). 5’-ACGGCCGCATCTTCTTGTGCA (forward primer for mus GAPDH), 5’-ACGGCCAAATCCGTTCACACC (reverse primer for mus GAPDH).

*Immunoblotting analysis.*

Colon were dissected longitudinally and washed with cold PBS. Sections of 3 cm distal to each colon or 293T cells were collected and mechanically homogenized with PRIA lysis buffer (Thermo Scientific) containing 1% PMSF according to the manufacturer’s protocol. The supernatant was collected and boiled in an SDS sample buffer for 10 min and analyzed by immunoblotting. Immunoblots were observed using Amey Imager 600RGB and quantified using ImageJ.

**Supplementary References**
