## Supplementary Figure1 for "ASB3 expression aggravates inflammatory bowel disease by targeting TRAF6 protein stability and affecting the intestinal microbiota"

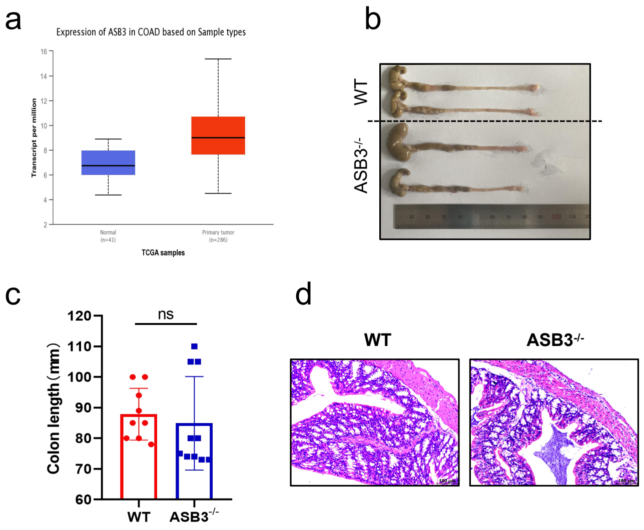


**Supplementary Fig. 1** (**a**) The transcription levels of ASB3 in colon adenocarcinoma (TCGA). (**b, c**) Colon length of WT and ASB3$^{-/-}$ mice treated without DSS. (**d**) Representative images of pathological H&E stained colon sections collected. Scale bar, 100 μm.
