## Supplementary Figure2 for "ASB3 expression aggravates inflammatory bowel disease by targeting TRAF6 protein stability and affecting the intestinal microbiota"

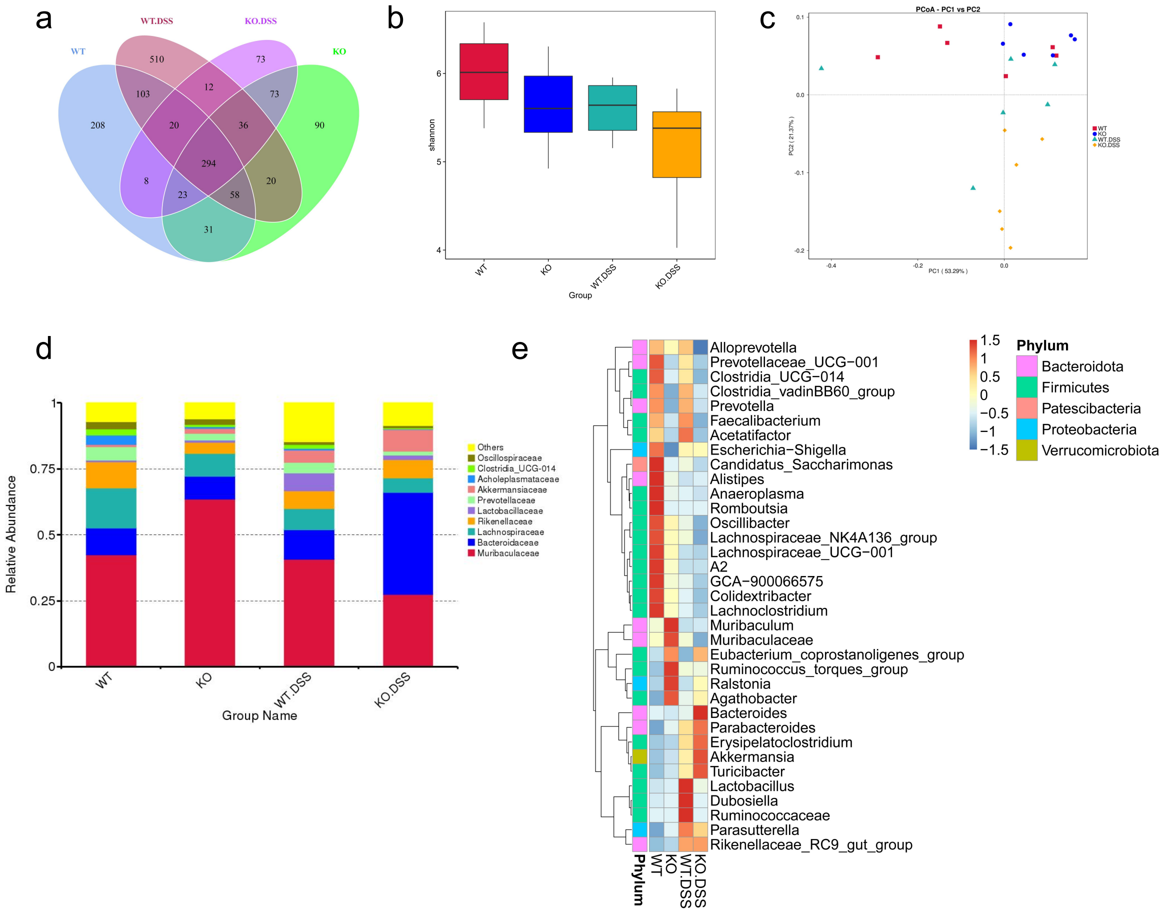


**Supplementary Fig. 2** (**a**) Venn diagram depicting the number of microbiota at the bacterial taxa (phylum, class, order and family) in the feces collected from WT and ASB3$^{-/-}$ mice at day 0 or day 6 of DSS-induced colitis. (**b**) Analysis of the bacterial diversity of fecal microorganisms from DSS-induced colitis in WT and ASB3$^{-/-}$ mice before and after. (**c**) Principal coordinate analysis (PCoA) of the weighted UniFrac distances of the fecal microbiota of WT and ASB3$^{-/-}$ mice pre- and post-DSS challenge (n=6 per group). (**d**) The relative abundance of microbial symbiont diversity was analyzed at the phylum level. (**e**) The relative abundance of microbial symbiont diversity was analyzed at the genus level.
