## Supplementary Figure3 for "ASB3 expression aggravates inflammatory bowel disease by targeting TRAF6 protein stability and affecting the intestinal microbiota"

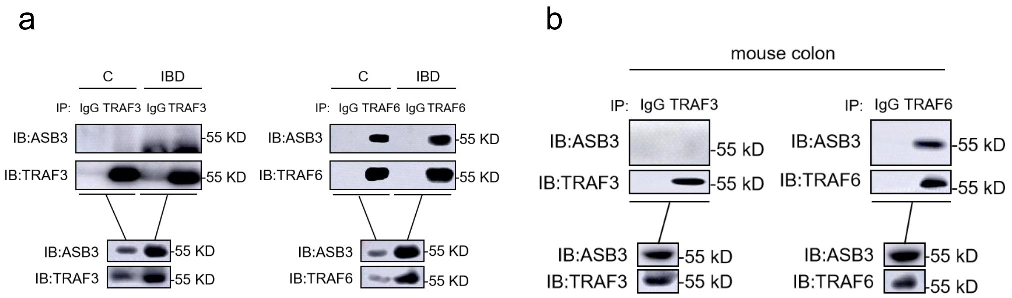


**Supplementary Fig. 3** (**a**) Total colonic tissue proteins from healthy and IBD samples. (**b**) Total colonic tissue proteins from DSS-treated mice. The samples were used in Co-IP assays with the indicated antibodies.
