## Supplementary Figure4 for "ASB3 expression aggravates inflammatory bowel disease by targeting TRAF6 protein stability and affecting the intestinal microbiota"

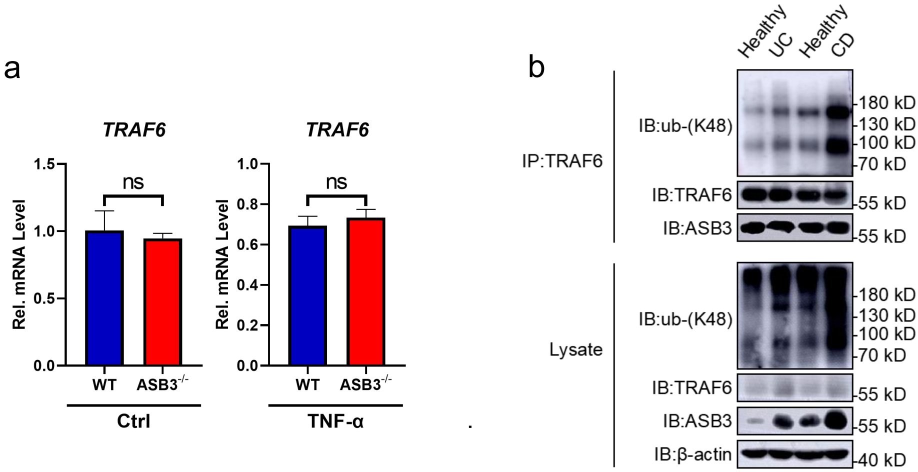


**Supplementary Fig. 4** (**a**) The mRNA expression levels of TRAF6 in organoids were determined by qPCR assay. (**b**) Total colonic tissue proteins from healthy and IBD samples. The samples were used in ubiquitination assays with the indicated antibodies.
