## Supplementary Figure5 for "ASB3 expression aggravates inflammatory bowel disease by targeting TRAF6 protein stability and affecting the intestinal microbiota"

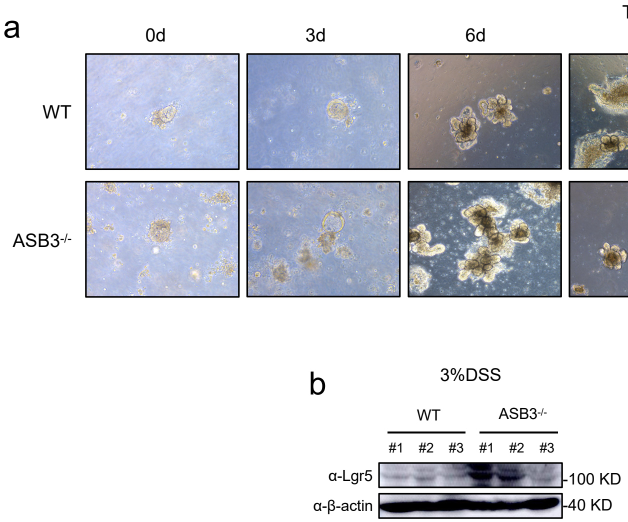


**Supplementary Fig. 5** (**a**) Intestinal stem cells were harvested from untreated WT and ASB3$^{-/-}$ mice, and the organoids were observed daily. Organoids were stimulated with mouse recombinant TNF-α for 24 h on day 7. (**b**) Western blotting was used to analyze Lgr5 protein expression in colons harvested from WT and ASB3$^{-/-}$ mice.
