## Supplementary Figure6 for "ASB3 expression aggravates inflammatory bowel disease by targeting TRAF6 protein stability and affecting the intestinal microbiota"

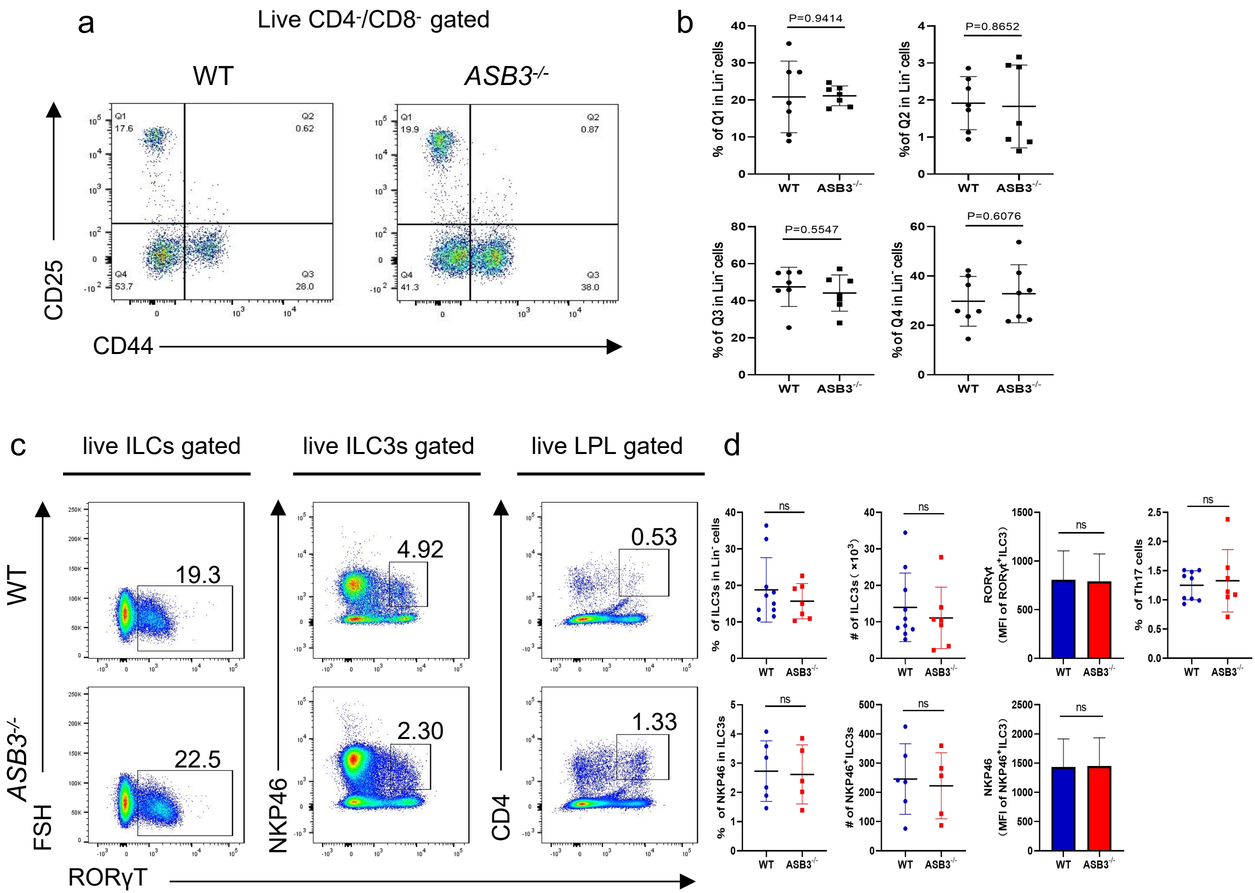


**Supplementary Fig. 6** (**a, b**) Frequencies of CD44-CD25+, CD44+CD25+, CD44+CD25- and CD44-CD25- T-cells in thymuses from WT and ASB3-/- mice. (**c, d**) Frequencies, absolute number and MFI of RORγT+ ILC3s, NKP46+ ILC3s and Th17 cells in cLPLs from WT and ASB3-/- mice.
